# Electron-bifurcation and fluoride efflux systems in *Acetobacterium* spp. drive defluorination of perfluorinated unsaturated carboxylic acids

**DOI:** 10.1101/2023.12.13.568471

**Authors:** Yaochun Yu, Fengjun Xu, Weiyang Zhao, Calvin Thoma, Shun Che, Jack E. Richman, Bosen Jin, Yiwen Zhu, Yue Xing, Lawrence Wackett, Yujie Men

**Affiliations:** Department of Chemical and Environmental Engineering, University of California, Riverside, Riverside, CA 92521, United States; Department of Environmental Chemistry, Swiss Federal Institute of Aquatic Science and Technology (Eawag), 8600 Dübendorf, Switzerland; Department of Biochemistry, Molecular Biology and Biophysics and Biotechnology Institute, University of Minnesota, Twin Cities, MN 55108, United States; SINOPEC Research Institute of Safety Engineering, Qingdao, 266071 Shandong, China

## Abstract

Enzymatic cleavage of C–F bonds in per- and polyfluoroalkyl substances (PFAS) is largely unknown but avidly sought to promote systems biology for PFAS bioremediation. Here, we report the reductive defluorination of α, β-unsaturated per- and polyfluorocarboxylic acids by *Acetobacterium* spp. Two critical molecular features in *Acetobacterium* species enabling reductive defluorination are (i) a functional fluoride efflux transporter (CrcB) and (ii) an electron-bifurcating caffeate reduction pathway (CarABCDE). The fluoride transporter was required for detoxification of released fluoride. Car enzymes were implicated in defluorination by the following evidence: (i) only *Acetobacterium* spp. with *car* genes catalyzed defluorination; (ii) caffeate and PFAS competed *in vivo*; (iii) models from the X-ray structure of the electron-bifurcating reductase (CarC) positioned the PFAS substrate optimally for reductive defluorination; (iv) products identified by ^19^F-NMR and high-resolution mass spectrometry were consistent with the model. Defluorination biomarkers identified here were found in wastewater treatment plant metagenomes on six continents.

## Main

Anthropogenic organofluorines, particularly per- and polyfluoroalkyl substances (PFAS), are among the most strictly regulated and concerning emerging contaminants^1,2^. The perfluorinated structure makes C–F bond dissociation energies high enough to evade oxidative, reductive and hydrolytic enzymes currently known^3,4^. Although C–F bond reduction is thermodynamically favorable overall^5^, the high energy (i.e., low reduction potential) required, together with fluoride toxicity and the short evolutionary time span of PFAS exposure to microbes, likely constrains PFAS biodefluorination^4^. Currently, PFAS defluorination has largely been shown to be indirectly triggered when weaker C–H or C–Cl linkages are present, which are more physiologically feasible for existing enzymes to attack^5–12^. Unstable intermediates such as *gem*-fluoroalcohols may be formed, leading to spontaneous C–F bond cleavage^6,7,13^. In contrast, direct enzymatic cleavage of C–F bonds in perfluorinated structures is rare^5^. For example, *Acidimicrobium* sp. A6 has been reported to defluorinate perfluorinated acids, but the underlying mechanisms and responsible enzyme(s) are yet to be identified^14–16^. Since bioremediation is considered to be a more cost-effective and environmentally friendly approach for *in-situ* cleanup of PFAS-impacted sites, it is vital to identify defluorinating microorganisms/enzymes and elucidate molecular mechanisms of C–F bond cleavage to allow systems biology and enzyme engineering to progress.

In previous studies, we observed reductive defluorination of an unsaturated perfluorinated compound, *E*-perfluoro-4-methylpent-2-enoic acid (PFMeUPA, (CF_3_)_2_CFCF=CFCOOH), by an anaerobic microbial community^17^. The defluorination was not catalyzed by reductive dehalogenases of the dechlorinating microorganisms in the community^17^. This led us to further identify microorganisms and enzymes responsible for C–F bond cleavage. Here, we report that specific *Acetobacterium* species, commonly occurring acetogens, can catalyze reductive defluorination of PFMeUPA. Defluorination occurring intracellularly accumulates toxic F^-^, thereby requiring functional F^-^ transporters. In the defluorinating *Acetobacterium* species, a heterodimeric Fluc F^-^ channel (CrcB_1_ and CrcB_2_) was employed for F^-^ detoxification. Evidenced by both *in vivo* experiments and *in silico* modeling, the electron-bifurcating caffeate reduction complex (CarABCDE)^18^ was involved in the reductive defluorination. Metagenomic screening of CarC homologs revealed a ubiquitous occurrence of potential defluorinating microorganisms in wastewater environments. Here, the identification of defluorinating pure cultures and the elucidation of molecular mechanisms of microbial reductive defluorination offer valuable insights into rational design and optimization of biocatalysts, which could be potentially applied for bioremediation of per- and polyfluorinated contaminants.

## Results

### PFMeUPA is reductively defluorinated at the *α*-C position by *Acetobacterium bakii*

We observed complete transformation of PFMeUPA (∼100 μM) within three weeks by *Acetobacterium bakii* grown on fructose (Figure 1a). The defluorination was significantly faster than what we reported previously (∼120 days) in an anaerobic mixed culture^17^. This indicated that PFMeUPA might only be biotransformed by a specific and minor microbial group within the community. Despite the different rates, *A. bakii* transformed PFMeUPA into the same intermediates and end products as the community with the same fluoride release at a near 1:1 molar ratio to the transformed PFMeUPA (Figure 1a). By comparing MS^2^ and ^19^F-NMR spectra of biological samples and isomeric products of chemical catalytic reduction of PFMeUPA, we confirmed the stereo-specific single isomer structure of the two microbial reductive defluorination transformation products (TPs): (i) the unsaturated form, (CF_3_)_2_CFCF=CHCOO^‒^, with an exact m/z of 256.9855 and designated “TP256” (the same nomenclature hereafter for other TPs) and (ii) its secondary hydrogenation product (TP259, (CF_3_)_2_CFCHFCH_2_COO^‒^) (Figure 1a, b and Figure S1-2). This demonstrates an α-C defluorination in the microbial system. Unlike a parallel abiotic reaction that we conducted here, only one of the separatable diastereomers from chemical synthesis was detected as a biohydrogenation product (TP276, (CF_3_)_2_CF(CHF)_2_COO^‒^) during PFMeUPA biotransformation by *A. bakii* (Figure 1a, b and Figure S3). The stereospecificity of the TPs demonstrates that those reactions carried out by *A. bakii* were enzyme mediated. Quantifying the two end TPs (TP259 and TP276) in the mixture by ^19^F-NMR spectra indicated that the majority (∼82%) of PFMeUPA was transformed via reductive defluorination (Figure 1b, c).

**Figure 1.**
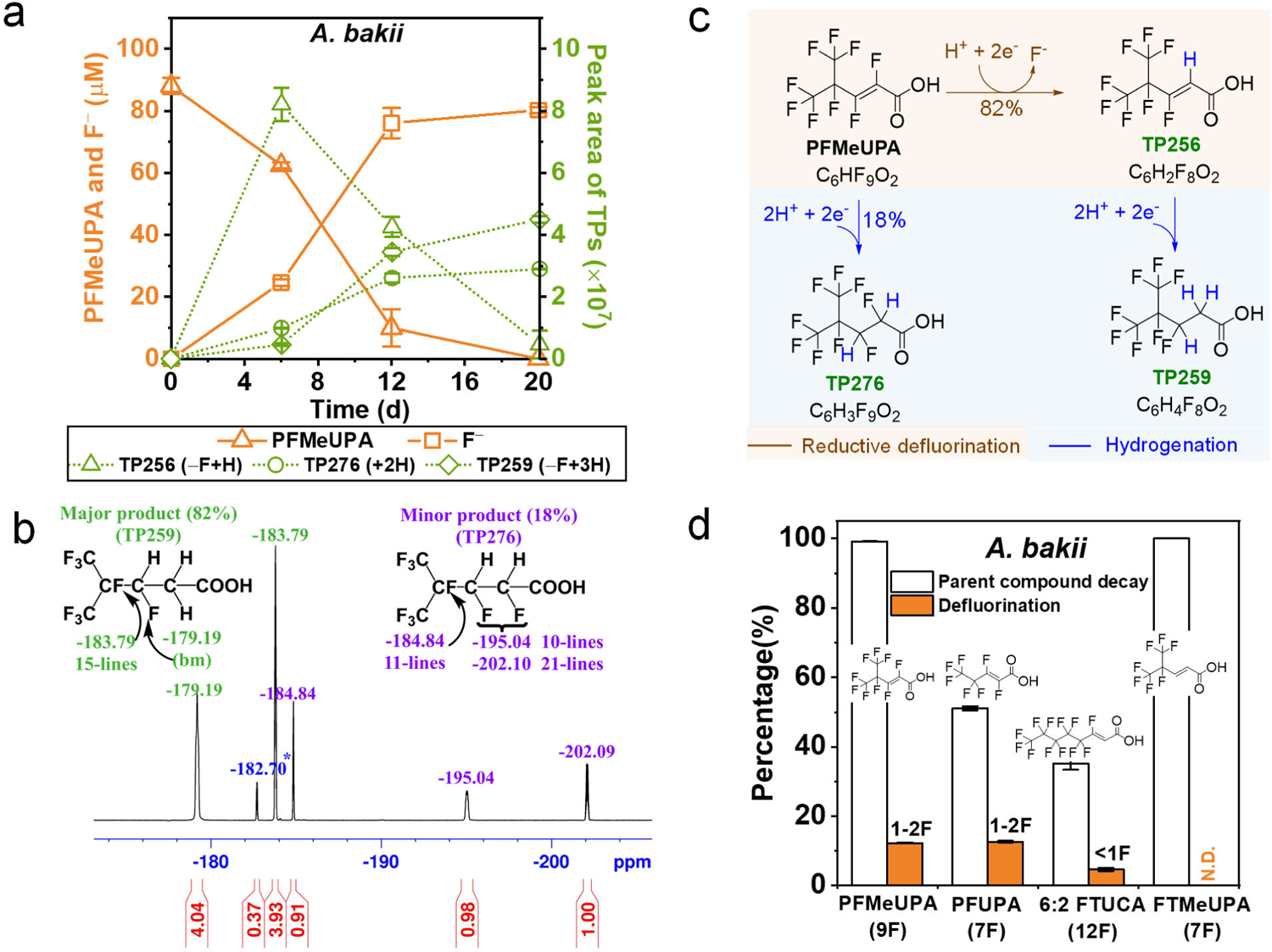
Defluorination of per- and polyfluorinated unsaturated carboxylic acids by *A. bakii*. **a.** PFMeUPA removal and the formation of fluoride and defluorination intermediate (TP256) and stable end products (TP276 and TP259); **b.** ^19^F-NMR spectra confirmation of defluorination intermediates and products (red numbers: integration of resonances; see Figure. S1‒S3, for the LC-HRMS/MS spectra); **c.** Confirmed biotransformation pathways of PFMeUPA; **d.** Substrate-specificity of *A. bakii* for reductive defluorination.

*A. bakii* also transformed two other unsaturated PFAS with structures similar to PFMeUPA, PFUPA (CF_3_CF_2_CF=CFCOOH) and 6:2 FTUCA (CF_3_(CF_2_)_4_CF=CHCOOH) (Figure 1d), via reductive defluorination and hydrogenation (Figure S4a-d). The total defluorination was similar to what we previously observed in an anaerobic microbial community^13^. While the community slightly defluorinated FTMeUPA ((CF_3_)_2_CFCH=CHCOOH)^17^, *A. bakii* only hydrogenated this structure without any F^‒^ formation (Figure 1d and Figure S4e-f). This indicated that while *A. bakii* carried out the reduction of a number of α/β unsaturated per- and polyfluorinated carboxylic acids, it only specifically cleaved C–F bonds on the unsaturated carbons. Thus, the previously reported defluorination of FTMeUPA at the tertiary C–F by the anaerobic community^17^ should be attributed to other pathways with different mechanisms.

### A functional F*^‒^* transporter is a prerequisite for intracellular defluorination

The reductive defluorination of PFMeUPA occurred intracellularly because we did not observe any defluorination activity in the cell-free spent medium of *A. bakii* grown on fructose (Figure S5). Given the cellular toxicity of F^‒^ at relevant concentrations, microorganisms capable of sustainable intracellular defluorination must possess detoxification mechanisms and actively export fluoride (F^‒^) ^19^. *A. bakii* possesses a heterodimer Fluc F^‒^ channel^19,20^ encoded by *crcB1 and crcB2*. Both genes were significantly (multivariate t-test, adjust *p* < 0.01) upregulated (∼ 2- fold change) in the presence of PFMeUPA compared to the non-spiked control (Figure S6d and Table S3). The expression of the Fluc channel in *A. bakii* is controlled by a transcriptional F^‒^ riboswitch located upstream of *crcB1* (Figure S7)^19,21^. The upregulation of *crcB* genes indicated the activation of fluoride detoxification by *A. bakii* in response to PFMeUPA defluorination and a functional fluoride exporter as one prerequisite for intracellular defluorination.

We then expanded the screening to five more *Acetobacterium* species (*A. bakii*, *A. woodii*, *A. paludosum*, *A. malicum*, and *A. fimetarium*) (Figure 2a). Interestingly, except for *A. fimetarium*, all the other strains exhibited defluorination activity (Figure 2a & b). In the genome of *A. fimetarium*, while *crcB1* is intact and homologous to that of *A. bakii*, *crcB2* was found to be truncated by 64 bp from the 3’ tail (Figure 2a). Given that both subunits must be present for a functional heterodimer Fluc F^‒^ channel^20^, we attributed *A. fimetarium*’s incapability of defluorination to its non-functional Fluc operon. To test this hypothesis, we introduced plasmid with *crcB1* and *crcB2* genes from five *Acetobacterium* species into *Escherichia coli* K-12 MG1655 *ΔcrcB* mutant strain, which is sensitive to F^‒^ and exhibited limited growth in the presence of 250 μM F^‒^. Except for *E. coli ΔcrcB* with *crcB1* and truncated *crcB2* from *A. fimetarium*, all other strains were rescued from F^‒^ toxicity (Figure 2c). Collectively, a functional F^‒^ exporter is essential for microorganisms to be capable of intracellular defluorination.

**Figure 2.**
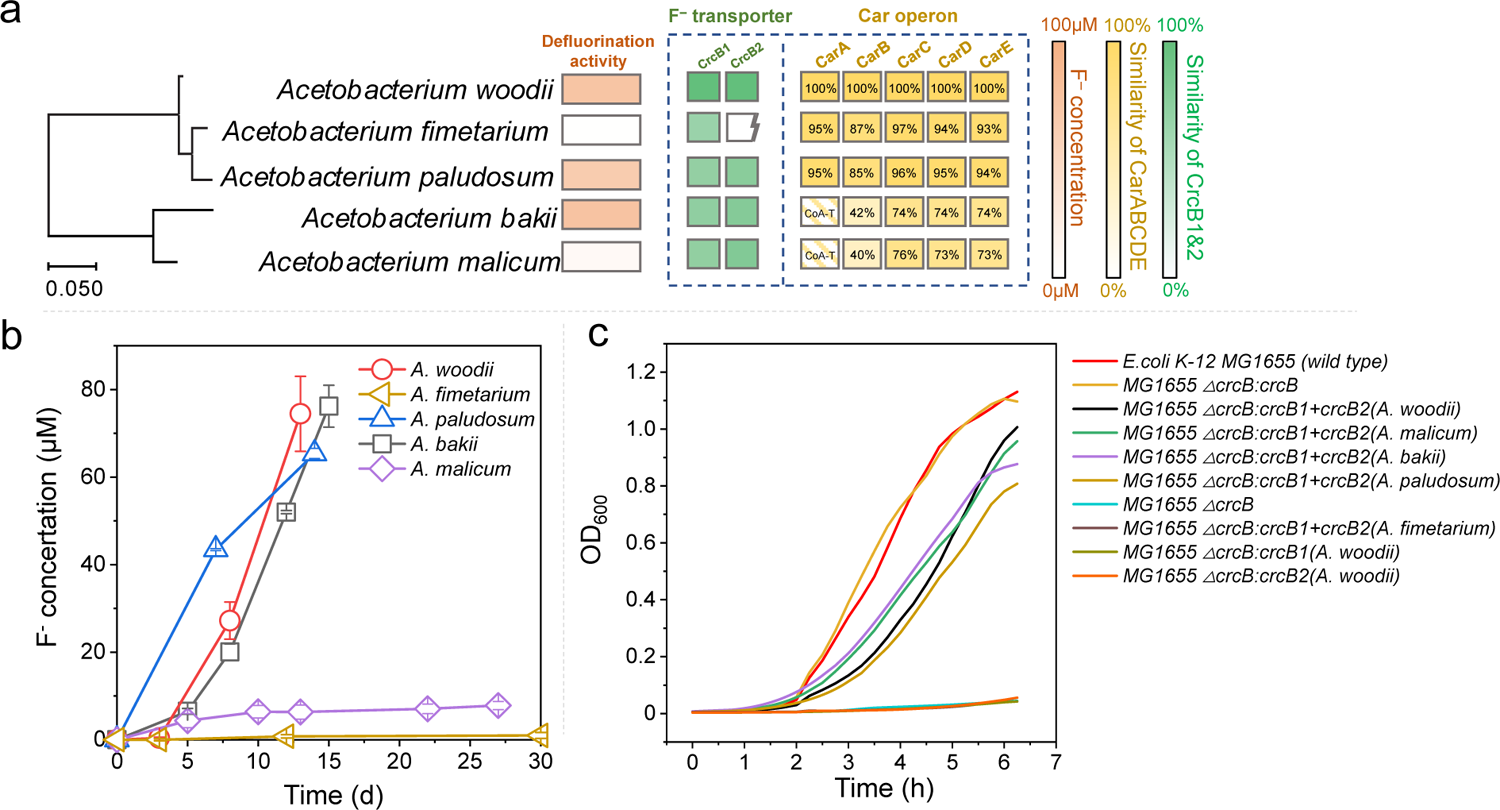
Comparison in PFMeUPA defluorination capability and fluoride resistance among *Acetobacterium* species. **a.** amino acid sequence similarities of the CrcB fluoride ion transporter and Car operon, CarABCDE among five *Acetobacterium* species and the corresponding total fluoride formation from ∼80 μM PFMeUPA; b. temporal fluoride formation from ∼80 μM PFMeUPA; c. growth restoration of *E. coli crcB*-knockout mutant, MG1655 Δ*crcB*, supplemented by *crcB* from different origins, including the native *crcB*, those from the five *Acetobacterium* species, and individual subunit genes, *crcB1* and *crcB2* from *A. woodii*.

### The electron-bifurcating caffeate reduction operon takes part in reductive defluorination

Besides the prerequisite of an active F^‒^ exporter for cell detoxification, we also explored potential enzymes involved in reductive defluorination. Based on the specificity of reductive defluorination to α/β-unsaturated per- and polyfluorinated carboxylic acids and the capability of *Acetobacterium* spp. in reducing caffeate, an α/β-unsaturated carboxylic acid, into hydrocaffeate^18^, we hypothesized that PFMeUPA was reduced by enzymes in the caffeate reduction pathway, which is encoded by the *car* operon (*carABCDE*) as previously characterized in *A. woodii*^22–24^.

We were not able to observe differences in the expression of *carABCDE* due to low levels of transcripts and proteins induced by the added PFMeUPA (Figure S6 and Table S1‒S3). Alternatively, we obtained strong evidence in the following aspects to demonstrate the involvement of the *car* operon in the reductive defluorination of PFMeUPA: (i) *in vivo* caffeate-PFMeUPA enzyme competition test; (ii) defluorination by additional *Acetobacterium* species with caffeate reduction activities; (iii) the consistency between the biochemical reactions observed for PFMeUPA and the pathways inferred based on the function of each gene in the *car* operon; (iv) *in silico* substrate-binding and modeling of CarC using the reported crystal structure^22^.

First, the addition of 5 mM caffeate in *A. bakii* completely inhibited the defluorination of PFMeUPA during the first week (Figure 3a) without significant inhibition on cell growth (Figure S8a). Defluorination activity was revived after caffeate was depleted after 12 days (Figure 3a and Figure S8b). PFMeUPA defluorination inhibited by caffeate at the early stage implied that caffeate competed with PFMeUPA for the binding site of the enzyme catalyzing reductive defluorination.

**Figure 3.**
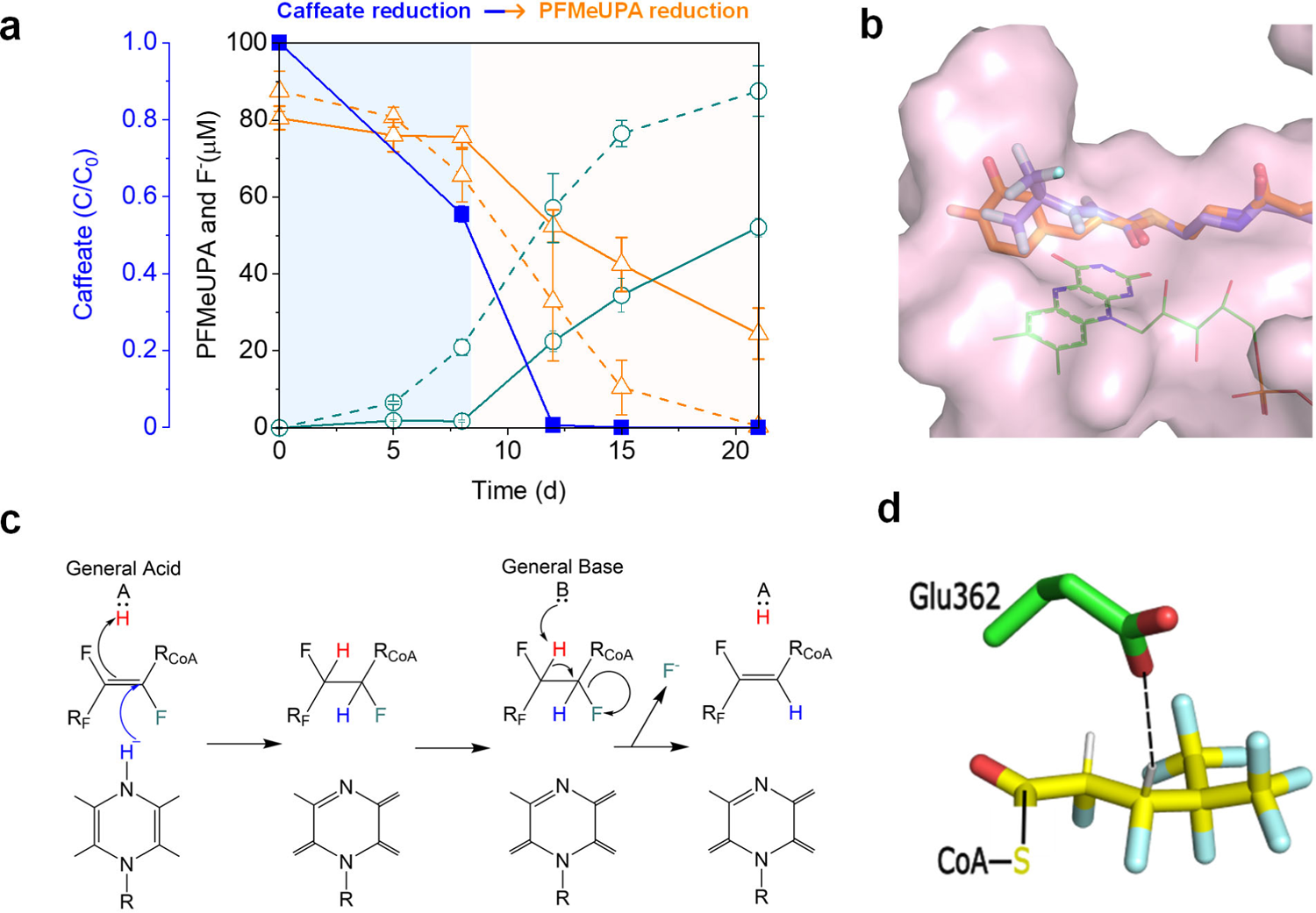
Reductive defluorination of PFMeUPA by enzymes involved in the caffeate reduction pathway. **a.** caffeate inhibition on PFMeUPA defluorination, orange: PFMeUPA, green: fluoride, blue: caffeate; solid line: with 5 mM caffeate; dashed line: without caffeate; b. docking of CoA thioesters of hydrocaffeic acid (orange) and PFMeUPA (purple) into CarC active site (pink) with the relative position of FAD shown (green); c. proposed reaction pathway showing reduction of fluorinated substrate by FAD (bottom) and elimination of fluoride; d. CoA thioester of PFMeUPA docked into CarC showing Glu362 ability to act as a general acid and base.

We then compared PFMeUPA defluorination activities of the four *Acetobacterium* species (i.e., *A. bakii*, *A. woodii*, *A. paludosum*, and *A. malicum*) with a functional Fluc F^‒^ channel and similar caffeate reduction operons (Figure 2a). Similar defluorination activities were observed for *A. bakii*, *A. woodii*, and *A. paludosum*, while *A. malicum* exhibited partial defluorination (Figure 2b).

Using the X-ray structure coordinates of the CarCDE enzyme complex^22^, we modeled the caffeoyl-CoA binding site of CarC to determine if caffeoyl-CoA and an analogous perfluorinated acyl-CoA could compete for binding and position the 2,3-enoyl double bond near the CarC flavin for reduction. Figure 3b shows the docking model of each CoA thioester in the surface cavity proposed for the X-ray structure of CarC to harbor caffeoyl-CoA,^22^ consistent with the structure of other enoyl-CoA reductases from *Clostridium* spp^25^. The modeling showed sufficient space in the binding site to accommodate the branched -CF(CF_3_)_2_ moiety in the same region as the catecholic aromatic ring of caffeoyl-CoA (Figure 3b). Most importantly, the double bond of each substrate is nearly superimposable with respect to the flavin cofactor previously identified to carry out the reduction reaction^22^. The modeling also identified a residue that is conserved across many enoyl-CoA reductases, Glu362, that was poised to within 2.2 Å of C-3 on caffeoyl-CoA or, as shown, C-3 of the analogous perfluorinated acyl thioester (Figure 3c, d). This immediately suggested a mechanism for defluorination in which Glu362 serves as a general acid to deliver a proton to the opposite face of the double bond as the hydride equivalent is delivered by the flavin cofactor to reduce the double bond (Figure 3c). After reduction, the deprotonated acid can then serve as a general base to stabilize removal of the proton from C-3 and facilitate the β-elimination of a fluoride anion from C-2, forming TP256. This enzyme-assisted defluorination explains the regiospecific fluoride elimination observed for the microbiological defluorination, whereas we see the elimination of fluorine from either C-3 or C-2, or both, in chemical reductions (Figure S1‒S3). This is consistent with the identification of TP259 as the sole product from an additional reduction of the double bond in TP256. Based on our modeling, no further defluorination is possible for TP256 because there is no fluorine at C-2 to undergo defluorination. The minor transformation product, TP276, would derive from the exit of the hydrogenation intermediate from the CarC active site before deprotonation occurred. This enzyme-assisted defluorination pathway (Figure S9) is also supported by the TP formation trend (Figure 1a), with TP256 first increasing then decreasing while both TP276 and TP259 increase over time. This argues against a precursor-product relationship between TP276 and TP256/TP259 that would be observed if defluorination were occurring freely in solution.

### Chain-length specificity and prevalence of microorganisms possessing Car operon (CarC) in nature

There are several microbial groups, including some *Clostridium* species, known to possess electron bifurcating enoyl-CoA reductases. We further tested *Clostridium homopropionicum* (DSM5847), a species that utilizes electron bifurcation in a fermentation pathway yielding propionic acid^26^. This pathway involves α, β-enoyl-CoA reduction reactions with substrates significantly shorter than caffeoyl-CoA. Analysis of the published genome sequence^27^ reveals gene constellations analogous to those in *Acetobacterium spp*., including homologs of CarCDE and fluoride ion export genes. Thus, *C. homopropionicum* is potentially capable of reductive defluorination, akin to *Acetobacterium spp.*, albeit with shorter-chain substrates. As expected, *C. homopropionicum* did not exhibit any transformation activity against PFMeUPA (Figure S10d). However, it did achieve slow defluorination (20 µM of F^-^ formation from 100 µM of parent compound over 15 days) for PFUPA (CF_3_CF_2_CF=CFCOOH) that is one carbon shorter than PFMeUPA (Figure S10). Considering that many other *Clostridium* spp. are known to catalyze the reduction of short-chain acyl-CoA substrates using genomically identifiable CarCDE homologs^28,29^, we think it is reasonable to suggest that other *Clostridia* could also react with short-chain unsaturated perfluorinated acids.

In light of the above, we collected sequences with >55% identity to *A. woodii* and *C. homopropionicum* from the non-redundant protein sequence database using the BLAST algorithm, created Hidden Markov Models (HMMs), and used the HMMs to search wastewater metagenomic sequences from the MGnify Database. Sequences with sample-origin metadata were used to create a map showing wastewater treatment plants containing moderate to high CarC sequence identity to those found in *Acetobacterium*, *Clostridium*, or both. In total, 723 metagenomes were identified with homologs, a breakdown by continent is shown in Table S4. In this context, wastewater treatment plants (WWTPs) have a significant probability of harboring microbes capable of degrading the type of PFAS studied here, and perhaps other fluorinated chemicals with α, β-unsaturation. This was validated by observing PFMeUPA defluorination activities in various environmental microcosms derived from not only WWTPs (i.e., activated sludge in anaerobic conditions and digester sludge) but also AFFF-impacted soil and groundwater communities (Figure S11).

## Discussion

The vast majority of organofluorines like PFAS are industrial chemicals, new to the biosphere, and among the most pervasive are perfluorinated acids^1,30–33^. Some consider these chemicals to be non-biodegradable given the high bond dissociation energy and low reduction potential of C–F bonds. Moreover, cytoplasmic defluorination, if occurred, would result in fluoride formation, which is highly toxic to cells. Here, we showed that combining a unique non-central metabolic trait (flavin-based electron bifurcation) and a co-opted detoxification trait (fluoride export) allowed *Acetobacterium* and *Clostridium* species to defluorinate unsaturated perfluorinated acids with different chain lengths.

Previous studies have demonstrated reductive defluorination of α, β-unsaturated perfluorocarboxylic acids by anaerobic microbial communities^17^, but common reductive dechlorinating bacteria and purified reductive dehalogenases did not react with fluorinated substrates^34^. Some biological reductions with the lowest reduction potential (< −500 mV) are carried out by flavin-based electron-bifurcating complexes^35,36^. These enzyme complexes are widespread in anaerobic Bacteria and Archaea where they couple ATP formation or utilization with the transformation of aryl/alkyl/alkenyl-CoA substrates, aldehydes, ketones, acids, hydrogen/protons and carbon monoxide^37^. The shared feature is the separation, or bifurcation, of electron pair energies such that one is boosted to a higher energy (lower potential) and the other to lower energy (higher potential)^35^. Here, we implicate one such system, CarCDE, in reductive defluorination. CarCDE is naturally evolved to reduce plant aromatic enoic acids such as caffeic acid, coupled with formation of reducing power for autotrophic growth or sodium transport to generate incremental ATP in energy-marginalized environments^38^. While there is little apparent structural relatedness to caffeic acid and PFMeUPA, modeling the respective CoA esters showed an ideal fit in the active site of CarC. This is also consistent with caffeate competition experiments implicating PFMeUPA as a possible electron acceptor alternative to caffeate and undergoing enzyme-assisted α-defluorination within the electron-bifurcating enzyme complex.

In different prokaryotes utilizing electron-bifurcation, the high energy electron reduces ferredoxins or flavodoxins that in turn reduce protons, dinitrogen gas, carboxylates to alcohols, benzenoid rings, and activate 2-hydroxyacyl-CoA dehydratases^35^. Indeed, the benzoyl-CoA reductase in *Thauera aromatica* has been shown to reductively defluorinate 4-fluoro-benzoyl-CoA via HF elimination and further reduction, a reaction analogous to that observed here with perfluoroalkenoic acids^39^. While this class I system uses ATP to drive the formation of high energy electrons, a class II benzoyl-CoA reductase uses two flavin-base electron bifurcation steps consecutively to catalyze benzene ring reduction^40^. This one-megadalton enzyme complex uses flavin, tungsten, selenium, and more than 50 iron-sulfur clusters to achieve the E^o’^ potential required for benzene ring reduction. These studies highlight an as-yet-unrealized prospect for endergonic reduction of a broader range of PFAS compounds by serving as the low-potential electron acceptor in the electron bifurcating system. It warrants future discovery or engineering of those enzyme complexes, overcoming the mechanistic/thermodynamic barriers, thus significantly shortening the time it could have taken by natural evolution.

A powerful enzyme complex capable of PFAS defluorination alone is insufficient due to fluoride toxicity. Intracellular fluoride release from obligately cytoplasmic defluorinating enzymes will immediately lead to fluoride binding to sensitive magnesium and calcium centers of enzymes, preventing cell growth^41^. It is necessary to couple C–F cleavage enzymes with rapid fluoride expulsion. Fluoride is abundant in the earth’s crust^42,43^. Thus, nature has evolved effective proteins to protect against fluoride leaking into cells from mineral fluoride^21,44^. Under fluoride stress, many native microorganisms express one of two families of membrane proteins to export F^−^, i.e., the CLC^F^ family of F^−^/H^+^ antiporters and the Fluc (CrcB) family of F^−^ exporting channels. CLC^F^ family proteins are homodimers, while Fluc family proteins include homodimers and heterodimers. Although the two types of F^−^ exporters differ in sequence, structure, and mechanism, they share the same physiological role in F^−^ detoxification. *A. fimetarium* with an intact Car operon but a non-functional CrcB was not able to defluorinate PFMeUPA at all. Such an incapability underscores the essential role F^−^ export proteins play in enabling cytoplasmic defluorination. This will be another critical design criterium, in addition to an active defluorinase, for efforts to bioengineer efficient PFAS-degrading microorganisms.

## Materials and Methods

### Fluorinated Compounds

Four fluorinated carboxylic acids were investigated based on their previously reported biodefluorination feasibilities in an anaerobic enrichment community.^13^ The chemicals were purchased from SynQuest Laboratories (Alachua, FL) and used without further purification. For all authentic standards, ten mM stock solutions were prepared anaerobically in autoclaved Milli-Q water (160 mL) sealed serum bottles and stored at room temperature until use. The limit of quantification for each standard compound was determined as the lowest concentration of calibration standards with a detection variation within ±20%. The detailed compound information is provided in Table S5.

The hydrogenation product of PFMeUPA was synthesized in-house using chemical catalysis. The catalyst, 39 mg of 5% Pd/C, and stir bar were charged to a Pyrex 6mL 14/20 RB flask, which was, in turn, positioned in the bottom of a 50mL Parr pressure reactor. The sealed reactor was evacuated (1.3 × 10^-4^ atm) and then filled with H_2_ gas. The reactor was maintained under a slow purge of H_2_ while 116 mg of PFMeUPA, followed by 2× washes totaling 2 mL of methyl-t-butyl ether (MTBE), was charged and magnetically stirred. The reactor was sealed and pressured to 3.74 atm with H_2_ and maintained for 4 hours. The reactor stirred overnight while the pressure was slowly released. Anhydrous HF was produced. After settling, the clear colorless supernatant was sampled (0.3 mL) and diluted to 0.7 mL with CDCl_3_ for the acquisition of ^19^F-NMR spectra using Varian Unity Inova 400MHz NMR Spectrometer for standard 5mm OD tubes. The products were also analyzed for structure confirmation by LC-HRMS/MS (as described below), and the LC chromatograph and MS^2^ fragmentation profiles (Fig. S1-S3) were used to compare with those of the products from biological reductive defluorination.

### Cultures and Growth Conditions

Five *Acetobacterium* and one *Clostridium* species were used to study the defluorination activities of the selected fluorinated compounds. *Acetobacterium bakii* (DSM 8239) was purchased from the American Type Culture Collection (ATCC); *Acetobacterium woodii* (DSM 1030), *Acetobacterium paludosum* (DSM 8237), *Acetobacterium malicum* (DSM 4132), *Acetobacterium fimetarium* (DSM 8238), and *Clostridium homopropionicum* (DSM 5847) were purchased from Deutsche Sammlung von Mikroorganismen und Zellkulturen (DSMZ). To test the biotransformation capabilities of individual species, we cultivated all cultures under their optimal growing conditions as instructed by DSMZ (Supplementary Methods).

### Biotransformation of Fluorinated Compounds by *Acetobacterium* spp and *Clostridium homopropionicum*

After the full growth of individual species, each culture was transferred (5%, v/v) into six serum bottles with 95 mL of fresh culture medium containing optimal primary substrates as suggested by DSMZ (*i.e.*, fructose for *A. woodii*, *A. paludosum*, *A. bakii*, *A. malicum*, *C. homopropionicum*; and lactate for *A. fimetarium*; see details in the Supplementary Methods). Two conditions were set up with triplicates: (1) growth positive control with no fluorinated compounds addition and (2) biotransformation group spiked with 50 – 90 μM of individual fluorinated compounds (i.e., one compound was spiked for each test). A triplicated heat-inactivated biomass adsorption control was also set up by inoculating 5 mL autoclaved (at 121 °C for 20 min for two cycles) culture into the same culture medium spiked with the same concentration of individual fluorinated compounds. Samples were taken subsequently during the incubation period for the measurement of the parent compound, transformation products (TPs), and F^-^. Briefly, at each sampling time, 3 mL aqueous suspension was centrifuged at 16,000 × g for 30 min (4 °C). Two mL supernatant was used for F^-^ measurement. The remaining (∼ 1 mL) was stored at 4 °C in the dark for subsequent LC-HRMS/MS measurement. The cell pellets from the biotransformation experiments were stored at –20 °C for DNA extraction.

### Differential gene expression by transcriptomic and proteomic analyses

Gene regulation at the transcriptional and translational level in *A. bakii* grown on 5g/L D-fructose with and without the addition of PFMeUPA (300 µM) was examined using RNA and protein sequencing. Details are included in the Supplementary Methods.

### DNA, RNA, and protein analyses

Detailed DNA extraction and quantitative polymerase chain reaction (PCR), RNA extraction, sequencing and bioinformatics analysis, and proteomics analysis are described in the Supplementary Methods.

### Functional validation of the fluoride ion transporter genes in *Acetobacterium* spp

The mutant strain *E. coli* K-12 MG1655 Δ*crcB* was constructed using the λ-red homologous recombination system and is sensitive to high fluoride concentration. The gene *crcB* from different *Acetobacterium* spp. was obtained using PCR. Purified PCR products were ligated to the linearized pBAD24 backbone. Recombinant plasmids were introduced into the mutant strain for rescue tests. Each strain was tested with LB medium supplemented with 250 µM of NaF. Cell growth was tracked via OD_600_ measurement. A detailed description of strain construction and rescue tests can be found in the Supplementary Methods.

### Substrate competition between caffeate and PFMeUPA in *A. bakii*

Five mM caffeate was added into six 160-mL serum bottles containing 95 mL culture medium and incubated overnight until caffeate was completely dissolved. Pre-grown *A. bakii* was then transferred (5%, v/v) into the above caffeate-containing medium bottles. Three of the six bottles were randomly picked as caffeate-only control and the remaining three were amended with ∼90 μM PFMeUPA. In addition, *A. bakii* (5%, v/v) was also inoculated into another bottle with growth medium only (without any addition of caffeate or PFMeUPA), serving as the positive control to confirm the normal growth activity. Samples were taken subsequently during the incubation period for the measurement of the parent compounds (i.e., PFMeUPA and caffeate), transformation products (TPs), and F^-^ as described below. Cells were collected and stored at –20 °C for subsequent DNA extractions and biomass growth measurements.

### Ultra-high performance liquid chromatography coupled to high-resolution tandem mass spectrometry (UHPLC-HRMS/MS) Analysis

For UHPLC analysis, 2 μL sample was loaded onto an Hypersil GOLD column (particle size 1.9 μm, 100 × 2.1 mm, Thermo Fisher Scientific) connected to an Hypersil GOLD guard column (particle size 3.0 μm, 10 × 2.1 mm, Thermo Fisher Scientific) and eluted with MilliQ water (A) and methanol (B) (both with 10 mM ammonium acetate) at a flow rate of 280 μL/min, with a gradient as follows: 95% A: 0 – 1 min, 95% – 5% A:1 – 6 min, 5% A: 6 – 8 min, and 95% A: 8 – 10 min. For HRMS/MS, mass full-scan was performed in the negative mode of electrospray ionization (ESI) at a resolution of 70,000 at m/z 200 and a scan range of m/z 50 – 750, and MS^2^ spectra were acquired in the same negative mode at a resolution of 14,000 at m/z 200 with a normalized collision energy of 25. Xcalibur 4.0 (Thermo Fisher Scientific) was used for data acquisition and analysis. Transformation products (TPs) screening was conducted using Compound Discoverer 3.1 (Thermo Fisher Scientific) as previously described.^6^ ChemDraw Professional 20.0 were used for drawing, displaying, and characterizing chemical structures.

### ^19^F and ^1^H NMR

^19^F- and ^1^H-NMR spectra on the biotransformation product of PFMeUPA by *A. bakii* were acquired on a Bruker Avance NEO 600-MHz NMR spectrometer with 5-mm H&F-C/N-D cryoprobe and adapter to 3-mm milli-tube. All spectra (^19^F and ^1^H) on products of PFMeUPA from abiotic catalytic hydrogenation were acquired on Varian Unity Inova 400MHz (proton) NMR spectrometer with standard 5 mm 4-nuclei (^1^H, ^31^P, ^13^C and ^19^F) probe.

### Fluoride ion measurement

The concentration of fluoride ion (F^-^) in the culture supernatant was measured by an ion-selective electrode (ISE, HACH, Loveland, CO) connected with an HQ30D Portable Multi Meter (HACH). The accuracy of the fluoride measurement by ISE was cross-validated using ion chromatography as described in our previous study.^17^ For the ISE measurement, a 100 μg fluoride ionic strength adjustment powder (HACH) was added into 2 mL culture supernatant, the F^−^ concentration was then measured by the ISE-Multi Meter system.

### Enzyme modelling

Models of the CarC active site were created by docking C_5_F_9_CO-CoA and caffeyol-CoA. The published crystal structure of *Acetobacterium woodii* CarC (PDB 6FAH) does not include caffeoyl-CoA in complex with the enzyme. Therefore, a highly similar CarC homolog from a Rat protein complexed with Acetoacetyl-CoA (PDB 1JQI) was aligned to the *Acetobacterium woodii* structure. After alignment, caffeoyl-CoA and C_5_F_9_CO-CoA were fitted to the electron density of the Acetoacetyl-CoA using Coot^45^. Restraint dictionaries for ligands were created on the Grade Web Server^46^. The protein-ligand complex structures were then subjected to geometric refinement using the Refmac5 tool within CCP4 with default parameters (except the the number of refinement cycles was set to 5 and the matrix weighting term was set to 0.0000001)^47^. Visualization was done in Pymol (The PyMOL Molecular Graphics System).

### Bioinformatics of CarC homologs in WWTPs metagenomes

Hidden Markov Models (HMM) were created to search a wastewater metagenome database for CarC homologs. These HMMs originated from two different seed sequences: Caffeyl-CoA reductase CarC (*Acetobacterium woodii* DSM 1030, NCBI AFA48354) and acyl-CoA dehydrogenase family protein (*Clostridium homopropionicum*, NCBI WP_052222365.1). For each seed, sequence databases with varying confidence levels were created using BLAST by gathering sequences at different percent identity thresholds from the non-redundant database. Proteins with 50% identity and higher have been reported to have less than 1 angstrom difference between their backbones^48^. Therefore, the lowest cutoff score was 55% identity for building a sequence library and constructing HMMs. Additionally, 74% identity was selected as the other cutoff, because that was the minimum similarity score identified in this study for other *Acetobacterium* CarC proteins capable of defluorination. These sequence databases were used for multiple alignments created with Clustal Omega^49^. Hidden Markov Models were generated from multiple alignments using hmmbuild version 3.1b2 (hmmer.org).

The newly created HMMs were used to identify CarC homologs in wastewater metagenomes and their locations were placed on the globalmap (Figure 4). A protein sequence database containing full-length cluster-representative proteins from wastewater was provided by MGnify from the May 2022 Protein database release^50^. MGnify database was searched using hmmsearch version 3.1b2. A sequence was considered a CarC homolog if the HMM score was higher than the lowest scoring sequence from which the database was built. Metadata for each sample was fetched using MgnifyR and samples without longitude and latitudes were removed^50^. Maps were built using ggplot2 in R^51^.

**Figure 4.**
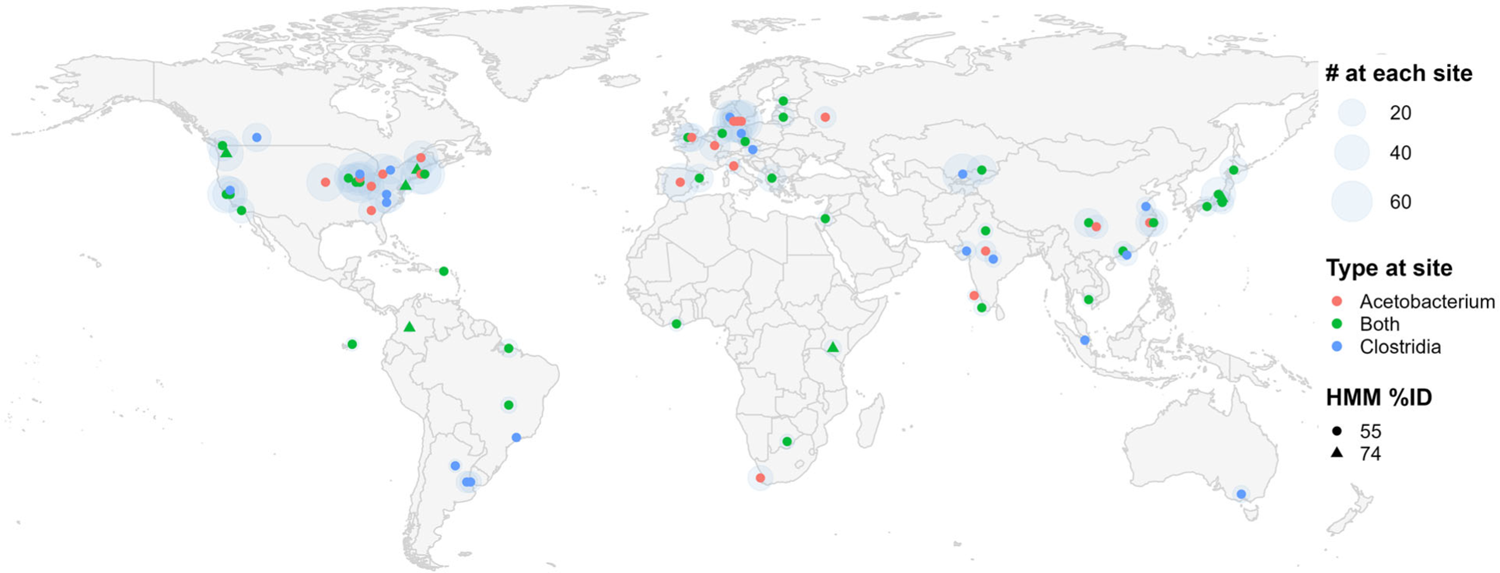
Presence of CarC homologs in WWTPs globally. The MGnify database was searched with Hidden Markov Models (HMMs) as described in the Methods section and generated using seed caffeoyl-CoA reductase (CarC) homologs from *Acetobacterium* and *Clostridium*. The search retrieved 723 metagenomes of which 499 had sufficient location metadata to plot the locations of the WWTP. In most instances, multiple metagenomic sequences were identified at a given site and the number is represented by the size of the blue circle. Some sites contained only *Acetobacterium*-type sequences, others only *Clostridium* type, and most contained both. Seven different WWTPs, shown as triangles, had a sample(s) that was identified with a higher confidence HMM (74% vs. 55%).

## Data availability

The sequencing raw reads of transcriptomes at two time points were deposited to the National Center for Biotechnology Information (NCBI) SRA database under accession No. PRJNA1029002 (20 day) and PRJNA1029168 (90 min). Preview links for reviewers: Long-term exposure data (https://dataview.ncbi.nlm.nih.gov/object/PRJNA1029002?reviewer=m05v8v797mgdodcricahaii4qm), and short-term exposure data (https://dataview.ncbi.nlm.nih.gov/object/PRJNA1029168?reviewer=59j0muutpovsoctnh853ehnk0h). The sequences will be made publicly available upon the publication of the associated research paper. The MGnify wastewater protein database used in this study is 9.3Gb in size. Due to this, interested parties may reach out to the authors to request the database.

## Supporting information

Supplementary Information for AceDef

Supplementary Tables for AceDef

## Acknowledgments

This study is supported by the Strategic Environmental Research and Development Program (ER20-1541 for Y.Y., B.J., W.Z., and Y.M.), the National Science Foundation (Award No. 1931941 for S.C.), and the National Institute of Environmental Health Sciences (Award No. R01ES032668 for S.C. and Y.M.), and partly by a grant from MnDRIVE Industry and the Environment Program (to L.P.W.).

## Author Contributions

Y.Y. and Y.M. conceptualized the research questions and hypothesis, designed the experiments and wrote the manuscript; Y.Y. and Y.M. selected the targeted acetogens species; Y.Y. conducted the biotranformation experiments and substrate competition experiments using *A. bakii*, *A. woodii*, and *A. fimetarium*; S.C., Y.Y., and W.Z, screened the defluorination capabilities of *A. malicum* and *A. paludosum*,; Y.Y. conducted the DNA and RNA extraction experiments and analyzed the whole-genome sequencing data and RNA sequencing data; Y.X. helped in establishing the RNA sequencing data analysis workflow and the results visualization; Y.Y., S.C., W.Z., and B.J. conducted the LC-HRMS/MS analysis; F.X. validated the functions of the fluoride ion efflux transporter in *Acetobacterium* spp.; Y.Z. constructed the PFMeUPA biodefluorination experiments using anaerobic activated sludge community and ground water community; C.T. constructed the CarC homologs analysis and modelling; J.E.R. performed the transformation product synthesis and ^19^F-NMR analysis. L.W. supervised the metagenomes data analysis, NMR analysis, and enzyme modeling; All authors reviewed and edited the manuscript; Y.M. supervised the entire project.

## Competing interests

The authors declare no competing financial interest.

## Notes

### Competing Interest Statement

The authors have declared no competing interest.

