## Supplementary Information for AceDef for "Electron-bifurcation and fluoride efflux systems in *Acetobacterium* spp. drive defluorination of perfluorinated unsaturated carboxylic acids"

### Table of contents

Supplementary Methods. .......... S3

Supplementary Tables. .......... S13

Supplementary Figures. .......... S13

Figure S1. Structure confirmation of TP256 by LC-HRMS/MS analysis. .......... S13

Figure S2. Structure confirmation of TP259 by LC-HRMS/MS analysis. .......... S14

Figure S3. The stereospecific formation of TP276 in PFMeUPA biotransformation according to LC-MS/MS analysis .......... S15

Figure S4. Biotransformation of PFUPA, 6:2 FTUCA, and FTMeUPA in *A. bakii*. .......... S16

Figure S5. Defluorination of PFMeUPA in *A. bakii*, heat-inactivated *A. bakii*, and cell-free spent medium of *A. bakii* grown on fructose .......... S17

Figure S6. Transcriptional and translational alterations of *A. bakii* during microbial defluorination of PFMeUPA. .......... S18

Figure S7. The *crcB* operon containing transcriptional $F^-$ riboswitch(es), *crcB1*, and *crcB2* in different *Acetobacterium* species. *A. woodii* has no $F^-$ riboswitch identified from its genome, *crcB2* in *A. fimetarium* is truncated. .......... S19

Figure S8. Cell growth of the *A. bakii* under different substrates exposure condition (a) and normalized fluoride concentration to cell growth (b). .......... S20

Figure S9. Schematic illustration of the metabolic pathways in *A. bakii*. .......... S21

Figure S10. Biotransformation and defluorination of a shorter chain unsaturated perfluorinated compound (PFUPA) by *Clostridium homopropionicum* DSM 5847. .......... S22

Figure S11. Biodefluorination of PFMeUPA in environmental microbial communities taken from different sites. .......... S23

**Supplementary Methods.**

***Five Acetobacterium species growth conditions.***

Five *Acetobacterium* and one *Clostridium* species were cultivated under growth conditions recommended by Deutsche Sammlung von Mikroorganismen und Zellkulturen (DSMZ). Specifically, *Acetobacterium bakii* was grown in DSMZ Medium 900 (pH 7.0), supplemented with 1 g/L yeast extract and 5 g/L fructose, and cultures were maintained at 20 °C. *Acetobacterium woodii* was cultivated at 30 °C in DSMZ Medium 135 (pH 8.0) containing 2 g/L yeast extract and 10g/L fructose as the primary substrate. *Acetobacterium malicum* was also grown at 30°C in DSMZ Medium 135 but with 1 g/L fructose as the primary substrate. *Acetobacterium paludosum* was kept at 20°C and grown in DSMZ Medium 614 (pH 7.0), which contains 0.5 g/L yeast extract and 10 g/L fructose. *Acetobacterium fimetarium* was cultivated at 30°C in DSMZ Medium 900 with 4 g/L lactate as the primary substrate. Lastly, *Clostridium homopropionicum* was incubated at 35 °C with DSMZ Medium 503 (pH 7.2), containing 2 g/L fructose as the primary substrate. The comprehensive list of Media for Microorganisms can be found in Media*Dive* (https://mediadive.dsmz.de/).

***Bacterial strains, plasmids, and primers.***

All genetically manipulated bacterial strains are derivatives of the wild-type strain *E. coli* K-12 MG1655. Strains and plasmids generated and used in this study were listed in Table M1 and Table M2, respectively. The *crcB* gene from different species was synthesized from the genomic DNA by polymerase chain reaction (PCR). Primers used in this study were provided in Table M3.

**Table M1.** Bacterial strains used and constructed in this study.

| Strains | Characteristics | Source or reference |
|---|---|---|
| *Escherichia coli* K-12 MG1655 | wild-type strain | [1] |
| | mutant strain △*crcB* | This study |
| | mutant strain △*crcB* with pBAD24-EC_crcB | This study |
| | mutant strain △*crcB* with pBAD24-AW_crcB1 | This study |
| | mutant strain △*crcB* with pBAD24-AW_crcB2 | This study |
| | mutant strain △*crcB* with pBAD24-AW_crcB | This study |
| | mutant strain △*crcB* with pBAD24-AM_crcB | This study |
| | mutant strain △*crcB* with pBAD24-AB_crcB | This study |
| | mutant strain △*crcB* with pBAD24-AP_crcB | This study |
| *Acetobacterium woodii* DSM 1030 | / | DSMZ |
| *Acetobacterium malicum* DSM 4132 | / | DSMZ |
| *Acetobacterium bakii* DSM 8239 | / | DSMZ |
| *Acetobacterium paludosum* DSM 8237 | / | DSMZ |
| *Acetobacterium fimaterium* DSM 8238 | / | DSMZ |

**Table M2.** Plasmids used this study.

| Plasmids | Characteristics | Source or reference |
|---|---|---|
| pKD4 | vector encoding *kanR*/*neoR* | Addgene[2] |
| pKM208 | λ-red encoding vector with a *lacZ* promotor | Addgene[3] |
| pBAD24-sfGFPx1 | pBAD24 carrying super fold GFP | Addgene[4] |
| pBAD24-EC_crcB | pBAD24 carrying *E. coli* K-12 MG1655 *crcB* | This study |
| pBAD24-AW_crcB1 | pBAD24 carrying *A. woodii crcB1* | This study |
| pBAD24-AW_crcB2 | pBAD24 carrying *A. woodii crcB2* | This study |

| | | |
|---|---|---|
| pBAD24-AW_crcB | pBAD24 carrying *A. woodii crcB1+crcB2* | This study |
| pBAD24-AM_crcB | pBAD24 carrying *A. fimaterium crcB* | This study |
| pBAD24-AB_crcB | pBAD24 carrying *A. bakii crcB* | This study |
| pBAD24-AP_crcB | pBAD24 carrying *A. paludosum crcB* | This study |
| pBAD24-AF_crcB | pBAD24 carrying *A. fimaterium crcB* | This study |
| pBAD24-SO_crcB | pBAD24 carrying *S. ovata crcB* | This study |

**Table M3.** Primers for *crcB1* and *crcB2* examinations.

| Primers | Sequence (5' - 3') | Target |
|---|---|---|
| KanR_F | ATTCGGCGTTACACTTATCACTCATACAAATCAAAT<br>AGCAGGATTTTGCAATGATTGAACAAGATGGA | *KanR* in pKD4 |
| KanR_R | ACGTCAGCAAGAATTCAAAACCCGCTTAATCAGCG<br>GGTTTTTTTTGGTCTTCAGAAGAACTCGTCAAG | |
| EC_crcB_F | GGGCTAGCAGGAGGAATTGTGTTACAACTTCTTTTA<br>GCAGTTTT | *crcB* in *E. coli* |
| EC_crcB_R | CGCCAAAACAGCCAAGCTTTAGTGTGCGGTTGAGG<br>CCG | |
| AW_crcB1_F | GGGCTAGCAGGAGGAATTATGCGAAAATACTATTT<br>CATTGCCG | *crcB1* in *A. woodii* |
| AW_crcB1_R | CGCCAAAACAGCCAAGCTTTAGTTGTAATCATAGCC<br>GAAGCG | |
| AW_crcB2_F | GGGCTAGCAGGAGGAATTATGATGGAAACTTTATG<br>TGTCATCC | *crcB2* in *A. woodii* |
| AW_crcB2_R | CGCCAAAACAGCCAAGCTTTAGAACGAGCTTGAGA<br>TTAAAAAACC | |
| AW_crcB_F | GGGCTAGCAGGAGGAATTATGCGAAAATACTATTT<br>CATTGCCG | *crcB1+2* in *A. woodii* |
| AW_crcB_R | CGCCAAAACAGCCAAGCTTTAGAACGAGCTTGAGA<br>TTAAAAAACC | |
| AF_crcB_F | GGGCTAGCAGGAGGAATTATGATGAAAAAATATTG<br>TTTTATTGCGGCC | *crcB1+2* in *A. fimetarium* |
| AF_crcB_R | CGCCAAAACAGCCAAGCTCTAATTGTAGTCAAAGC<br>CGAACTGGG | |
| AM_crcB_F | GGGCTAGCAGGAGGAATTATGAAAAAATATTATTT<br>TATCGCCGC | *crcB1+2* in *A. malicum* |

| | | |
|---|---|---|
| AM_crcB_R | CGCCAAAACAGCCAAGCTTTAACCAATTATTGCGCATGAAATCAG | |
| AB_crcB_F | GGGCTAGCAGGAGGAATTATGCGAAAGTATTATTTCATTGCGGC | *crcB1+2* in *A. bakii* |
| AB_crcB_R | CGCCAAAACAGCCAAGCTTTAATGGATTATTAAAATTGAAACAAAGTT | |
| AP_crcB_F | GGGCTAGCAGGAGGAATTATGAAAAAGTATTATTTTATTGCTGCCG | *crcB1+2* in *A. paludosum* |
| AP_crcB_R | CGCCAAAACAGCCAAGCTTCAGTGCGGAGCGATGATAAAA | |

Note: bases with underlined are homology arms, and bases in bold come from two sides of target genes.

***Gene knockout of crcB in E. coli K-12 MG1655***

The λ-red homologous recombination system was applied for *E. coli* gene knockout. The plasmid pKM208 encoding the recombinase was chemically transformed into competent *E. coli* MG1655. The expression was induced by adding isopropyl β-D-1-thiogalactopyranoside (IPTG) to a final concentration of 1 mM. The aminoglycoside 3'-phosphotransferase (kan) gene *kanR* with 50 bp homology arms of *crcB* on each side was amplified from the plasmid pKD4 by PCR using primers KanR_F and KanR_R (Table M3). To introduce purified *kanR* inserts into electrocompetent cells, electroporation was conducted on MicroPulser$^{TM}$ Electroporation Apparatus (Bio-Red) by using the default mode "Ec2" under the manufacturer's instructions. After electric shock, bacteria were mixed with LB medium and cultured at 30°C and 180 rpm for 1 h. Cells were then centrifuged, resuspended, and plated on LB agar medium containing 30 μg·mL$^{-1}$ kanamycin. After culturing overnight, colonies on the plate were picked and verified by PCR. Colonies with successful gene knockout were cultured at 37°C to eliminate pKM208.

***Recombinant plasmid construction and transformation***

To obtain the linearized vector for ligation, the plasmid pBAD24-sfGFPx1 bought from Addgene was digested by restriction enzymes EcoRI and HindIII. Inserts were obtained by PCR amplification using PrimeSTAR Max DNA Polymerase (TaKaRa Bio USA Inc.). PCR components included template (1 μL), the forward primer (0.5 mM, 0.25 μL), the reverse primer (0.5 mM, 0.25 μL), 2× PrimeSTAR Max Premix (25 μL), DMSO (0.5 μL), and 23 μL of $H_2O$. The program was set as the following: 1 cycle of initial denaturing at 98°C for 3 min, 30 cycles of denaturing at 98°C for 10 s, annealing at 56°C for 5 s, elongation at 72°C for 7 s, and a final extension at 72°C for 3 min. PCR products were examined by electrophoresis in 1% (w/v) agarose gels in 1× TAE buffer, stained with 10000× Biotium GelRed Nucleic Acid Gel Stain (Thermo Fisher Scientific), and visualized by Gel Doc™ EZ System (Bio-Red). After purification by Purelink™ PCR Purification Kit (Thermo Fisher Scientific), PCR products were ligated to the linearized pBAD24 by In-Fusion Snap Assembly cloning kits (TaKaRa Bio USA Inc.), under the manufacturer's instructions. Successful ligation was verified by agarose gel electrophoresis.

The ligation mixture was chemically transformed into competent *E. coli* DH5α. After growing overnight on LB plates supplemented with 50 $\mu g \cdot mL^{-1}$ ampicillin, transformants were picked, cultured, and screened for correct plasmids. PCR and Sanger sequencing were applied to verify gene integrity. Recombinant plasmids were extracted using QIAprep Spin Miniprep Kit (QIAGEN) and transformed into the chemically competent mutant strain *E. coli* K-12 MG1655 Δ*crcB* followed by the same procedure for selection.

***Rescue of E. coli K-12 MG1655 ΔcrcB from $F^-$ toxicity***

Unlike the wild-type strain, the *crcB* knockout (KO) mutant, *E. coli* K-12 MG1655 Δ*crcB*, is sensitive to high fluoride concentration (Figure M1). For the rescue test, the fluoride concentration was set to be 250 μM that could greatly inhibit the growth of the mutant strain.

Colonies of each strain were inoculated into fresh LB medium and cultured at 37°C and 180 rpm. To maintain plasmids, 50 μg·mL$^{-1}$ ampicillin was added. When $OD_{600}$ reached 0.3 to 0.5, 2 μL of culture was inoculated into 198 μL of LB medium supplemented with 250 μM sodium fluoride. Bacteria were cultured in a 96-well plate at 37°C in a microplate reader (BioTek Instruments, Inc.), and $OD_{600}$ was measured every 15 min. Neither Arabinose or glucose was added for the induction or the repression of gene expression, because the background expression level of *crcB* was sufficient for protection.

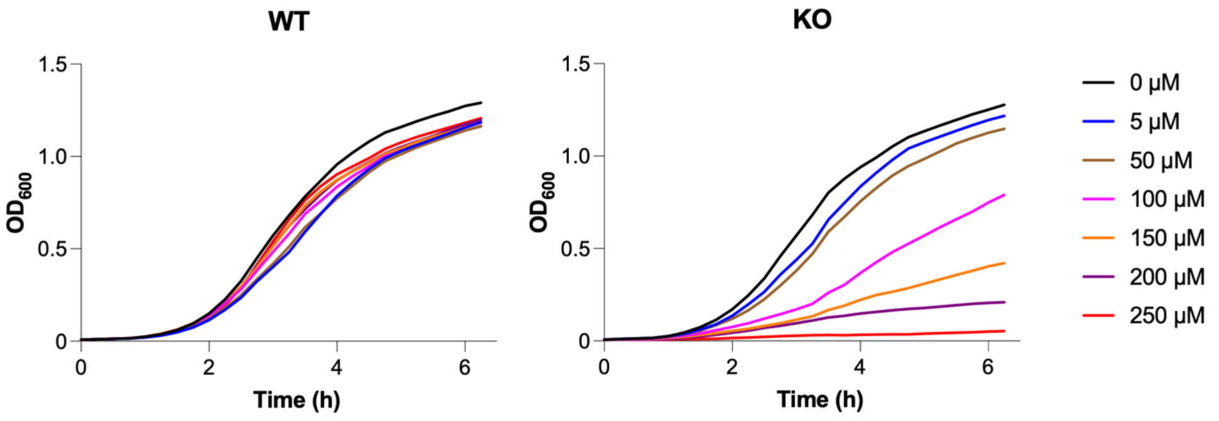


**Figure M1.** Growth curves of the wild-type (left) and the Δ*crcB* KO mutant (right) of *E. coli* K-12 MG1655 with different fluoride concentrations in the medium.

***DNA Extraction of A. bakii and Quantitative Polymerase Chain Reaction.***

Biomass from 1.5 mL of the culture was sampled from each biological replicate. Genomic DNA was extracted using a DNeasy PowerSoil Kit (QIAGEN, Germantown, MD) according to the manufacturer's instructions. Cell growth was measured by quantitative PCR (qPCR) using primers targeting the 16S rRNA genes of *A. bakii* (forward: 5'-GAGTACGACCGCAAGGTTGA-3', reverse: 5'-ATGCACCACCTGTCACTC TG-3'). The primers were designed and validated using Primer-BLAST (https://www.ncbi.nlm.nih.gov/tools/primer-blast/) and synthesized by Integrated DNA

Technology (IDT). Genomic DNA was quantified by NanoDrop One (Thermo Fisher Scientific) and served as the qPCR standards. PowerUp SYBR Green reagents (Thermo Fisher Scientific) were used for qPCR according to the manufacturer's instructions as reported in our previous study.[1] Briefly, the 20 µL qPCR reaction mixture contained 2.5 µL of the gDNA sample or genomic DNA standard, 10 µL of 2× PowerUp SYBR Green master mix solution, and 1.25 µL of 10 µM forward and reverse primers. The PCR procedure included an initial deactivation at 95 °C for 2 min, followed by 40 thermal cycles at 95 °C for 1 s and then at 60 °C for 30 s.

***Differential Gene Expression in A. bakii with the addition of PFMeUPA.***

As described in the previous biotransformation experimental section, after the growth of *A. bakii* has been observed, the culture was transferred (5%, v/v) into six serum bottles amended with 95 mL fresh medium with all essential nutrients. The freshly transferred cultures were first cultivated under room temperature for 30 h. Once the growth of the microorganisms was observed, 300 µM of PFMeUPA was then added to three out of the six bottles, while the other three were kept as non-PFMeUPA exposure controls. To investigate the short-term and long-term differential gene expression between PFMeUPA and non-PFMeUPA exposure groups, 9 mL culture was taken at 90-min and 20-days after adding the PFMeUPA. The sample was centrifuged at 16,000 × g for 30 min (4 °C). The supernatant was used for $F^-$ and LC-HRMS/MS measurement. The cell pellets were stored at –80 °C. For the 90-min samples, due to the low biomass availability, all of the cell pellets collected from the 9 mL culture were used for RNA extraction. For the 20-day samples, 3 mL of the cells were used for RNA extraction and the remaining 6 mL cells were sent to University of California, Los Angeles Proteome Research Center (UCLA-PRC) for proteomics analysis.

***RNA Extraction and RNA Sequencing.***

RNA was extracted using acid-phenol: chloroform: isoamyl alcohol (25: 24: 1) and precipitated in ethanol at – 20 °C as previously described[2]. RNA was cleaned up using the RNeasy PowerClean Pro CleanUp Kit (QIAGEN) according to the manufacturer's instructions. Contaminating DNA in the RNA samples was removed by Turbo DNase Kit (Thermo Fisher Scientific) following the manufacturer's instructions. The quality of RNA was examined by agarose gel electrophoresis. The RNA samples were sent to MiGS for RNA sequencing (RNA-seq). Briefly, the samples were first treated with Invitrogen DNAse (RNAse free) to remove any DNA contaminants. The cleaned RNA samples were then subjected to a library preparation using Illumina's Stranded Total RNA Prep Ligation with Ribo-Zero Plus kit and 10bp IDT for Illumina indices. For the 20-day samples, the sequencing was done on a NextSeq2000 giving 1 × 75 bp reads. For the 90-min samples, the sequencing was performed on a NextSeq2000 with 2 × 50 bp reads output. The demultiplexing, quality control, and adapter trimming of raw reads were performed by MiGS with bcl-convert (v3.9.3).

***Bioinformatics of the RNA sequencing Data.***

Differential gene expression analyses were performed on the Department of Energy Systems Biology Knowledgebase (KBase) platform (https://www.kbase.us/)[3]. Briefly, the RNA-seq reads from individual controls (i.e., with and without PFMeUPA) were uploaded to Kbase and grouped into one RNA-seq sample set. The sequencing reads were then aligned to the annotated reference genome (i.e., *A. bakii*) using HISAT2 v2.1.0[4]. The aligned RNA-seq were subjected to StringTie v1.3.3b for transcripts assembly[5, 6]. The normalized abundance of each transcript expressed in FPKM and TPM as Expression Matrix objects were provided by StringTie, and the  calculate counts-based differential gene expression under the two conditions were analyzed using DESeq2 v1.20.[7]. Genes were considered to have significantly differential expressions based on the following criteria: (i) FDR adjusted p-value $<0.05$; (ii) $>$ 2-fold difference in either TPM or FPKM values between the two exposure conditions.

***Proteomics Analysis***

Protein extraction and digestion, peptide analysis and protein identification from the 6 mL samples were conducted by UCLA-PRC. For sample preparation, bacteria pellets were lysed in lysis solution (8M Urea, 0.1M Tris-HCl pH 8.0, 1%SDS) containing AEBSF, pepstatin, leupeptin and benzonase. Samples were incubated with rotation for 30 min at temperature and then sonicated for 10 seconds in an ice-water bath using an ultrasonic cell homogenizer. Lysates were clarified by centrifugation at 16,100g for 15 min at 4°C. Protein amounts in each Lysate were normalized by absorbance at 280 nm and aliquoted, 50 µg, for the following digestion. For digestion and desalting, 50 µg protein aliquots were reduced and alkylated via sequential 20 min incubations with 5 mM tris(2-carboxyethyl)phosphine (TCEP) and 10 mM iodoacetamide at room temperature in the dark while being mixed at 1200 rpm in an Eppendorf thermomixer. Ten µl of carboxylate-modified magnetic beads (CMMB and also widely known as SP31) were added to each sample. Ethanol was added to a concentration of 50% to induce protein binding to CMMB. Then, the CMMB were washed 3 times with 80% ethanol and resuspended with 50µl of 50mM Tetraethylammonium bromide (TEAB). The obtained protein was digested overnight with 0.1 µg LysC (Promega) and 0.8 µg trypsin (Pierce) at 37 °C. Following digestion, 1ml of acetonitrile was added to each sample to increase the final acetonitrile concentration to over 95% and induce peptide binding to CMMB. The CMMB were washed 3 times with acetonitrile and the peptide was eluted with 50 µl of 2% DMSO. Eluted peptide samples were dried by vacuum centrifugation and reconstituted in 5% formic acid before analysis by LC-MS/MS. For the LC-MS acquisition and analysis, peptide samples were separated on a 75µM ID, 25cm $C_{18}$ column packed with 1.9 μM $C_{18}$ particles (Dr. Maisch GmbH HPLC) using a 140-minute gradient of increasing acetonitrile concentration and injected into a Thermo Orbitrap-Fusion Lumos Tribrid mass spectrometer. MS/MS spectra were acquired using Data Dependent Acquisition (DDA) mode. MS/MS database searching was performed using MaxQuant (1.6.10.43) against the Xanthobacter autotrophicus reference proteome from Uniprot (UP000036873, *Acetobacterium bakii* DSM 8239, 3495 entries). Statistical analysis of MaxQuant label-free quantitation data was performed with the artMS Bioconductor package, which performs the relative quantification of protein abundance using the MSstats Bioconductor package (default parameters). Intensities were normalized across samples by median-centering the $\log_2$-transformed $MS^1$ intensity distributions. The abundance of proteins missing from one condition but found in more than 2 biological replicates of the other condition for any given comparison were estimated by imputing intensity values from the lowest observed MS1-intensity across samples and p-values were randomly assigned to those between 0.05 and 0.01 for illustration purposes. To further address the missing data in the protein intensity dataset through a statistic approach, data produced from the MaxQuant were analyzed in RStudio (version 4.2.2) using a "missForest" package, which predicted and imputed all missing values using random forest algorithm and returned a completed data matrix. The PCA plot shown in Figure S6 was done using the imputed data matrix.

**Supplementary Tables.**

All supporting tables are uploaded separately in an associated Excel file (named: *SupportingTables_AceDef.*).

**Supplementary Figures.**

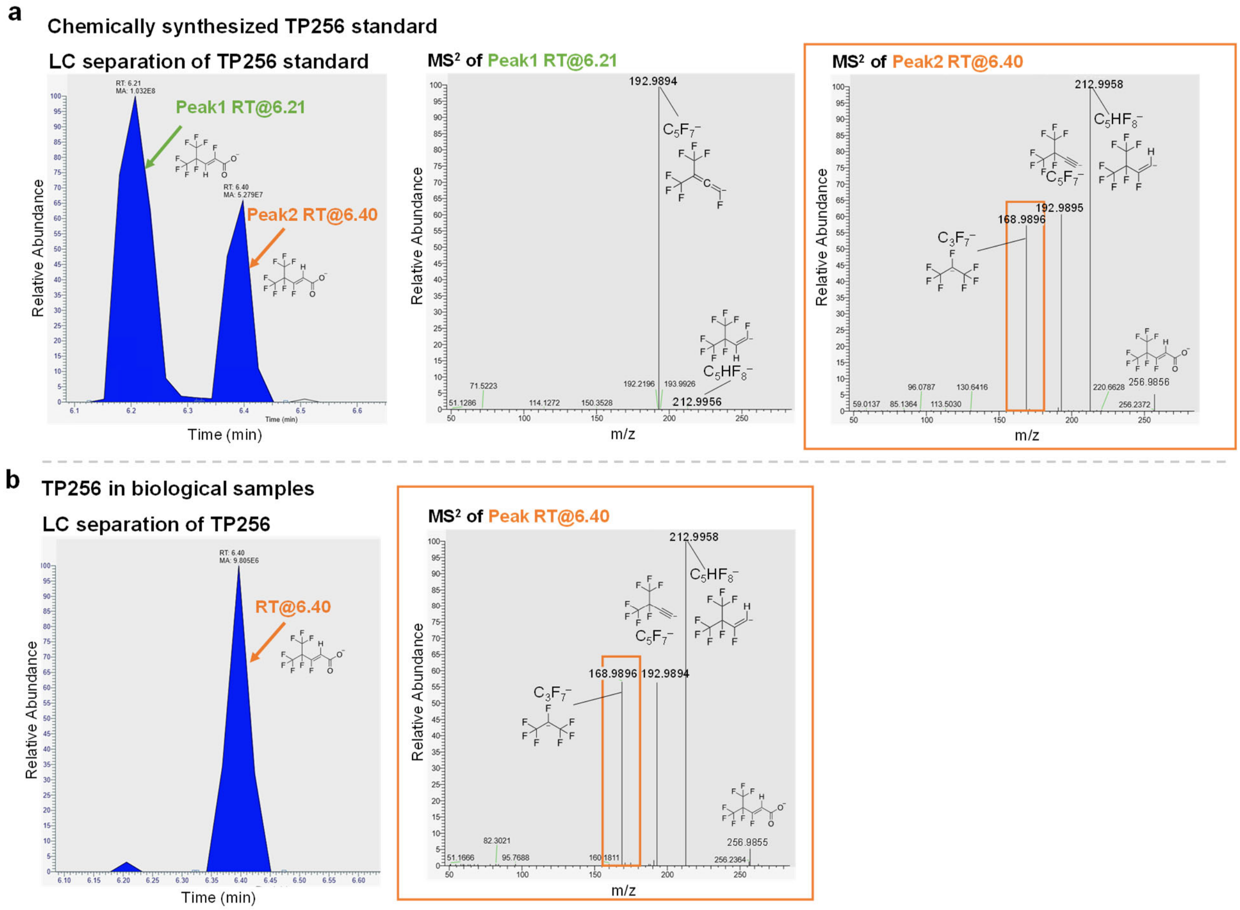


**Figure S1**. **Structure confirmation of TP256 by LC-HRMS/MS analysis.** a, LC chromatograph (left) and $MS^2$ fragmentation spectra (middle and right) of two TP256 isomers (with the same m/z = 256.9855 and formula $C_6HF_8O_2^-$) formed from chemical reduction of PFMeUPA. b, LC chromatograph (left) and $MS^2$ fragmentation spectra (right) of the nearly exclusively formed TP256 isomer in PFMeUPA biotransformation samples. The microbially formed TP256 was confirmed to have the same structure as an α-defluorination product (Peak 2) by comparing the retention time (RT), signature $MS^2$ fragment (i.e., $C_5F_7^-$ as a single dominant fragment for the β-defluorination product (Peak 1) and $C_3F_7^-$ as a unique fragment for the α-defluorination product (Peak 2) in Panel a), as well as the relative intensities of fragments. Note: The first three digits of the exact m/z were used to name the TPs, e.g., TP256 refers to the peak with an m/z = 256.9855.

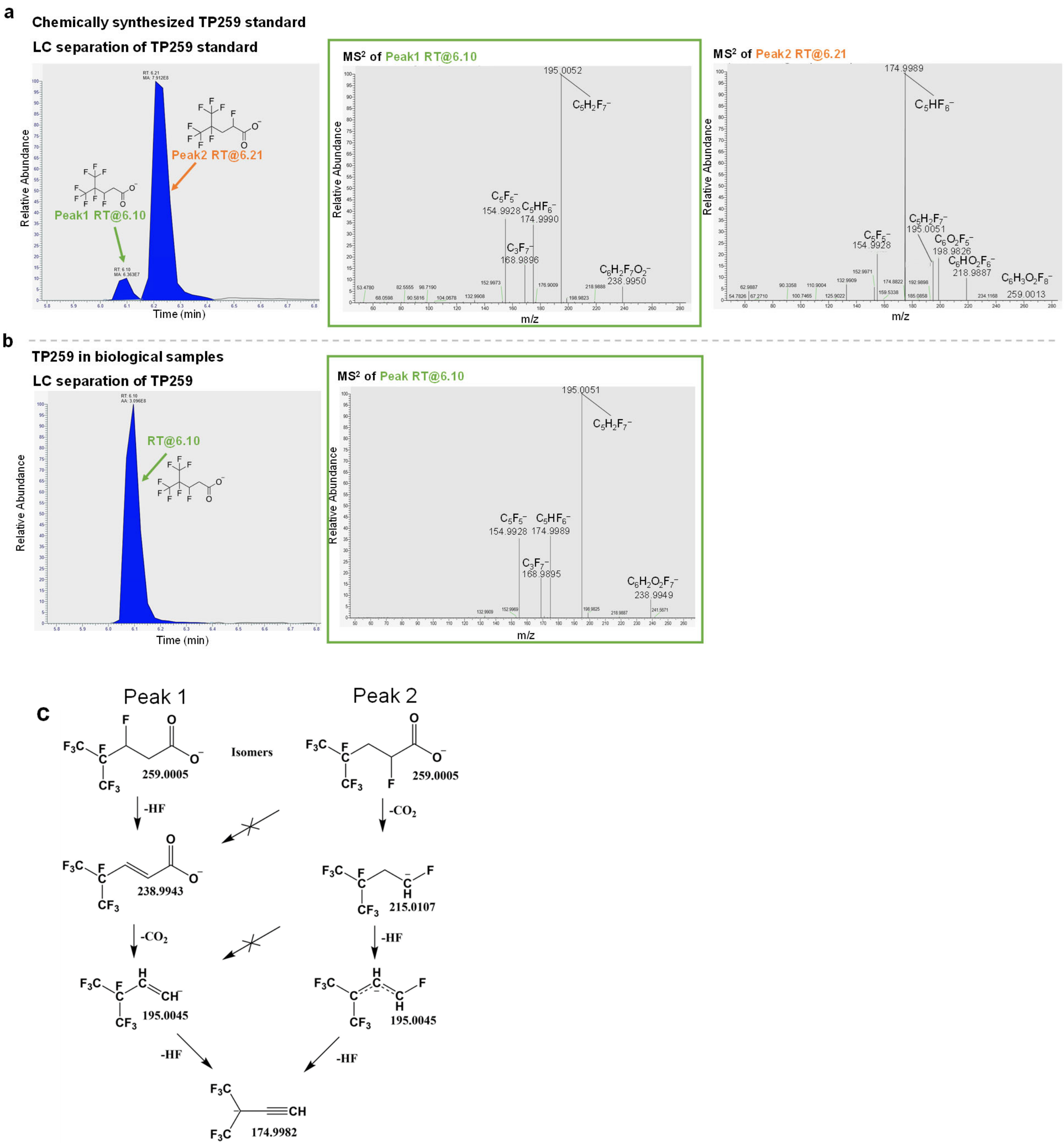


**Figure S2**. **Structure confirmation of TP259 by LC-HRMS/MS analysis.** a, LC chromatograph (left) and MS$^2$ fragmentation spectra (middle and right) of two TP259 isomers (with the same m/z = 259.0005 and formula $C_6H_2F_8O_2^-$) formed from chemical reduction of PFMeUPA. b, LC chromatograph (left) and MS$^2$ fragmentation spectra (right) of the exclusively formed TP259 isomer in PFMeUPA biotransformation samples. c, the interpretation of MS$^2$ fragments of the two isomers of TP259 (theoretical m/z values are indicated with a mass error < 5ppm from the detected ones). By comparing the retention time (RT) and MS$^2$ fragments, we confirmed the structure of microbially formed TP259 to be the same as Peak 1, the α-defluorinated isomer.

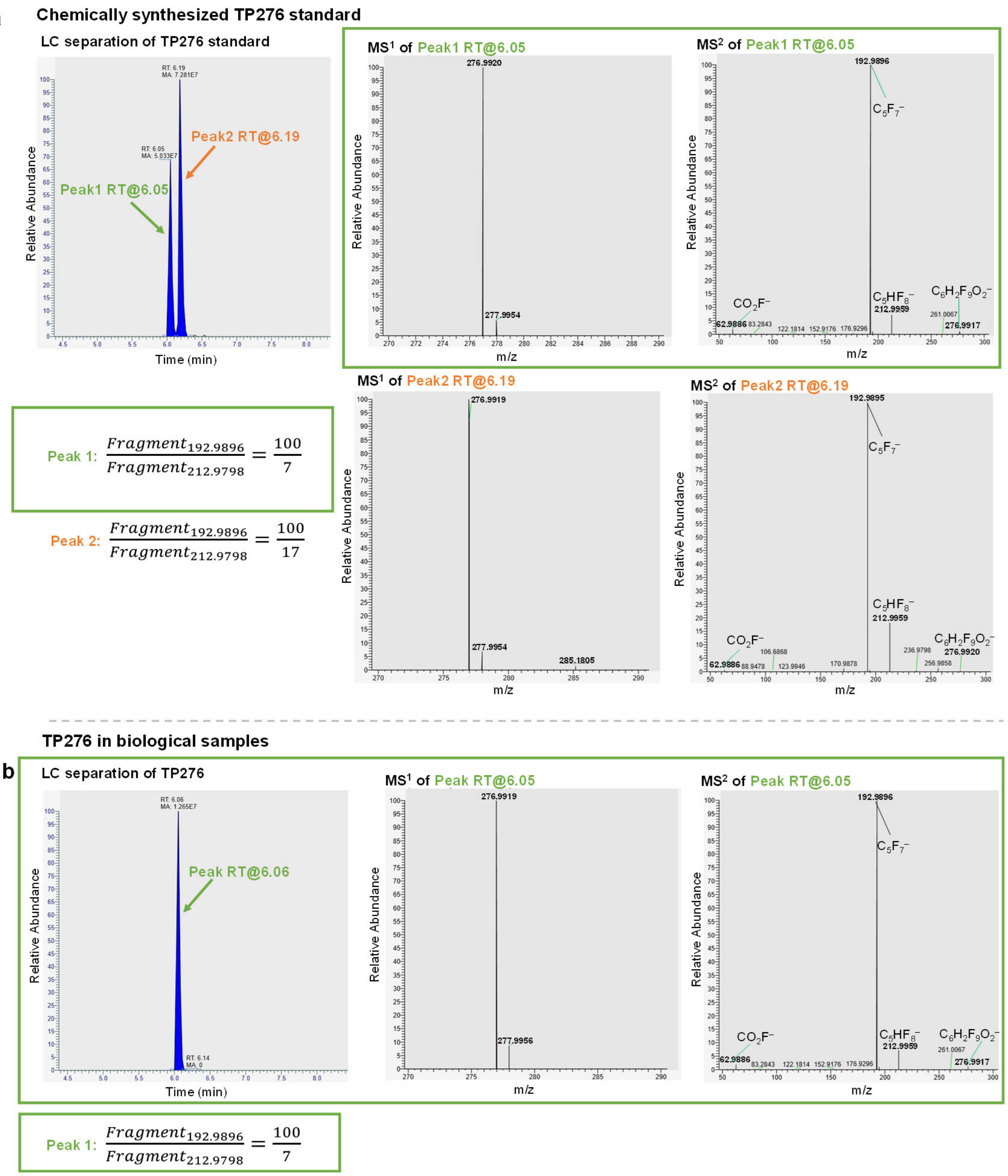


**Figure S3**. **The stereospecific formation of TP276 in PFMeUPA biotransformation according to LC-MS/MS analysis.** a, the LC chromatograph (left), MS full-scan (middle), and $MS^2$ fragmentation spectra (right) of the two diastereomers of TP276 from chemical reduction of PFMeUPA. b, the LC chromatograph (left), MS full scan (middle), and $MS^2$ fragmentation spectra (right) of the single diastereomer of TP276 from PFMeUPA biotransformation. By comparing the retention time (RT) and the ratio of two dominant MS2 fragments (m/z = 192.9896 and 212.9798) we confirmed the structure of TP276 is the same as Peak 1 in panel a but were not able to differentiate the two structures represented by Peak 1 &2 due to the same fragment patterns.

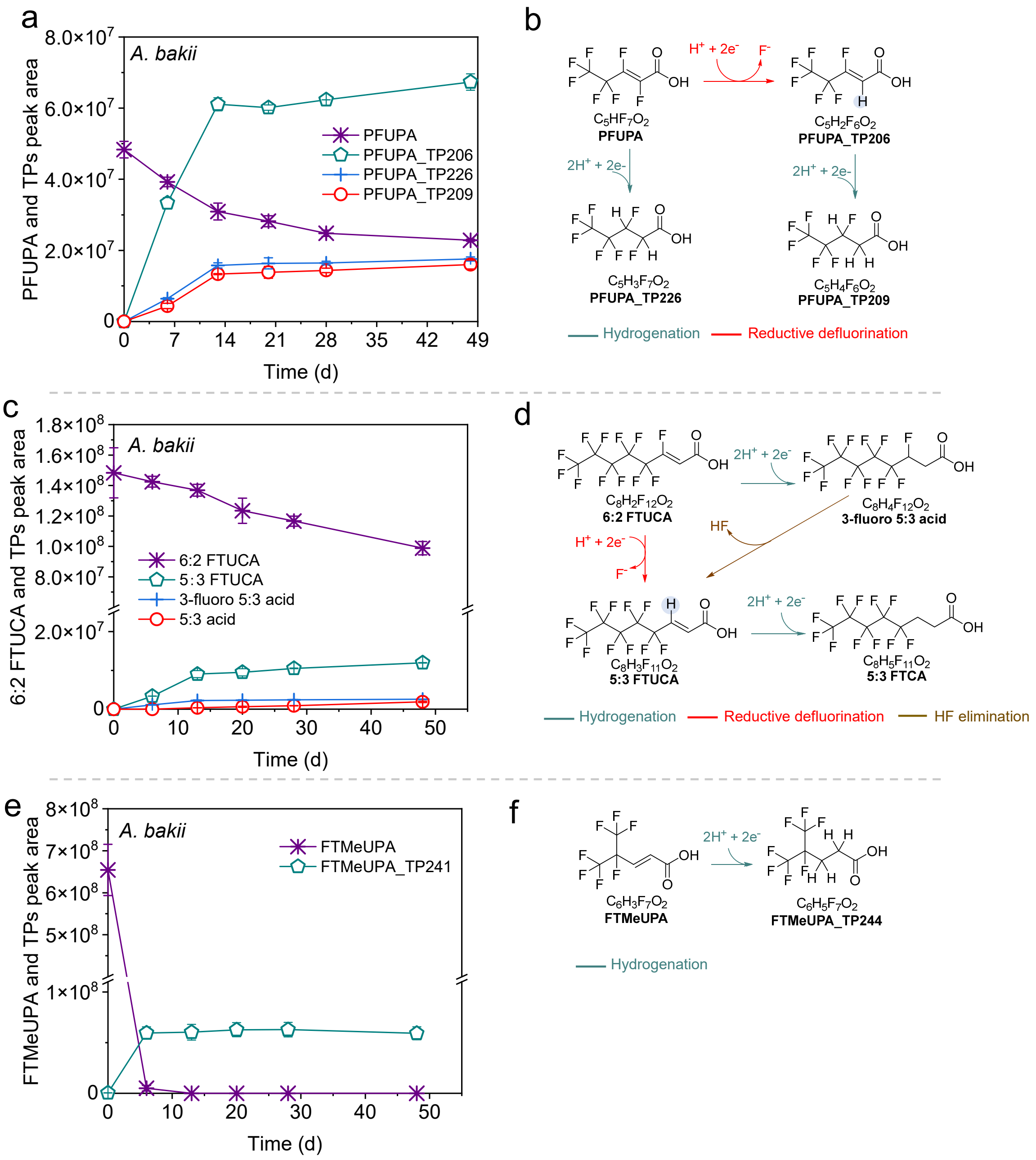


**Figure S4**. **Biotransformation of PFUPA, 6:2 FTUCA, and FTMeUPA in *A. bakii*.** a, Biotransformation of PFUPA and TPs formation. b, Biotransformation pathways of PFUPA. c, Biotransformation of 6:2 FTUCA and TPs formation. d, Biotransformation pathways of 6:2 FTUCA. e, Biotransformation of FTMeUPA and TP formation. f, Biotransformation pathway of FTMeUPA. (Note: the first three digits of the exact m/z were used to name the TPs).

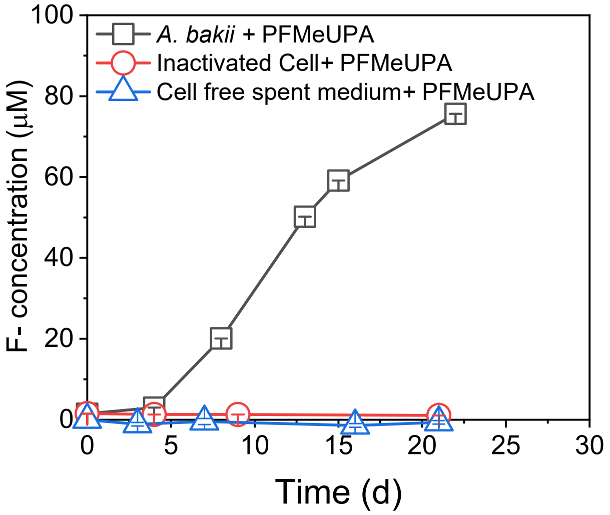


**Figure S5. Defluorination of PFMeUPA in *A. bakii*, heat-inactivated *A. bakii*, and cell-free spent medium of *A. bakii* grown on fructose.**

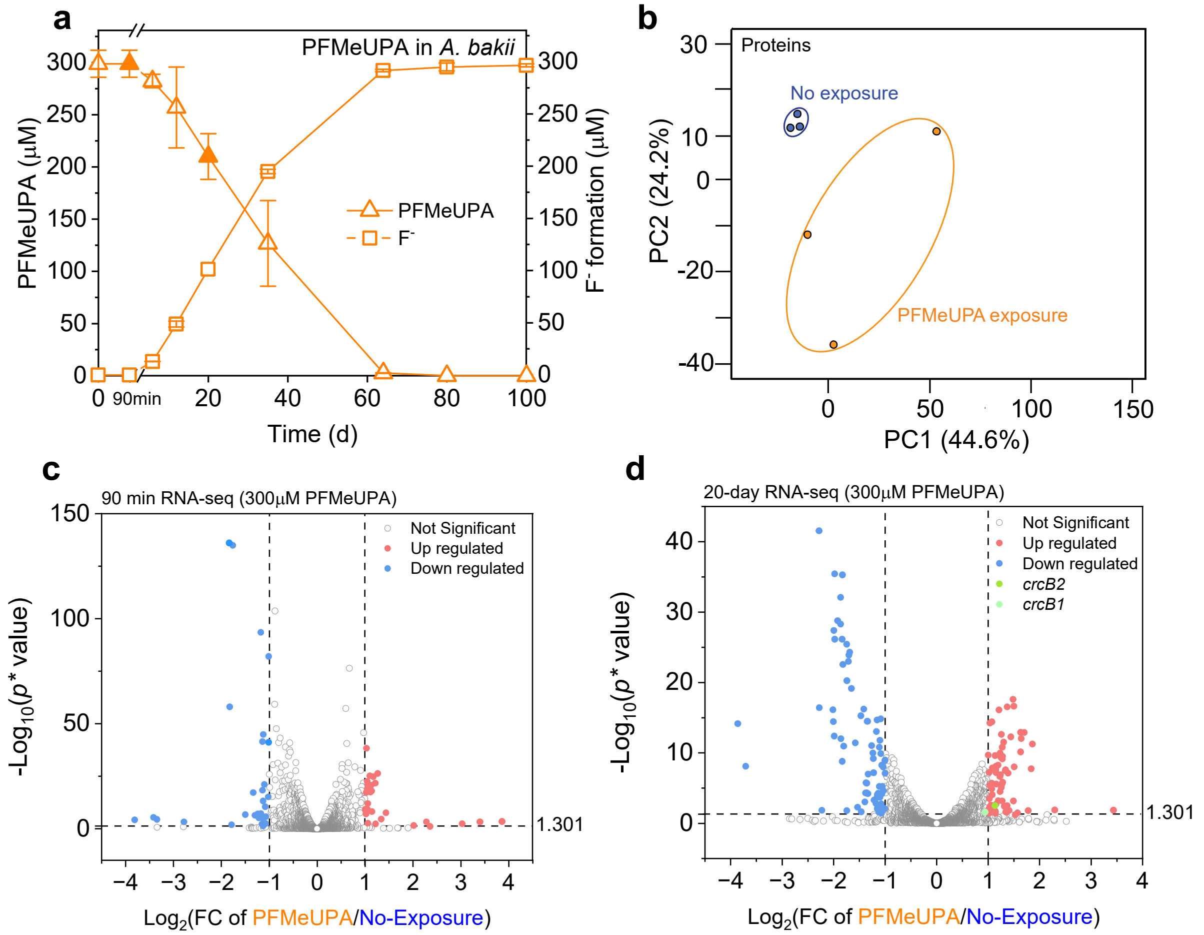


**Figure S6. Transcriptional and translational alterations of *A. bakii* during microbial defluorination of PFMeUPA.** a, 300 µM PFMeUPA degradation and $F^-$ formation (solid symbols indicate samples were taken for RNA and/or Protein analysis). b, Principal component analysis of *A. bakii* protein intensities under PFMeUPA-exposed and non-PFMeUPA exposed conditions (samples were taken at 20-day). c, d, Volcano plots of *A. bakii* transcriptomes under short-term (90 min after adding PFMeUPA) and long-term (20-day after adding PFMeUPA) PFMeUPA exposure conditions (|log2 fold change of PFMeUPA/No-Exposure| > 1, adjusted p-value < 0.05, significance limits are marked with dashed lines). The detailed differential gene expression analysis results are included in Tables S1 (proteome), Table S2 and S3 (RNA-seq).

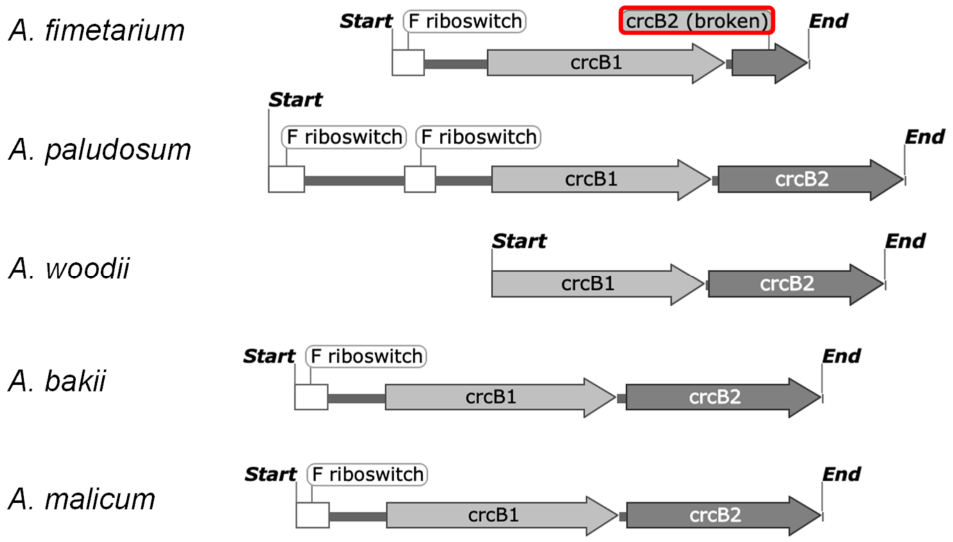


**Figure S7. The *crcB* operon containing transcriptional F⁻ riboswitch(es), *crcB1*, and *crcB2* in different *Acetobacterium* species.** *A. woodii* has no F⁻ riboswitch identified from its genome, *crcB2* in *A. fimetarium* is truncated.

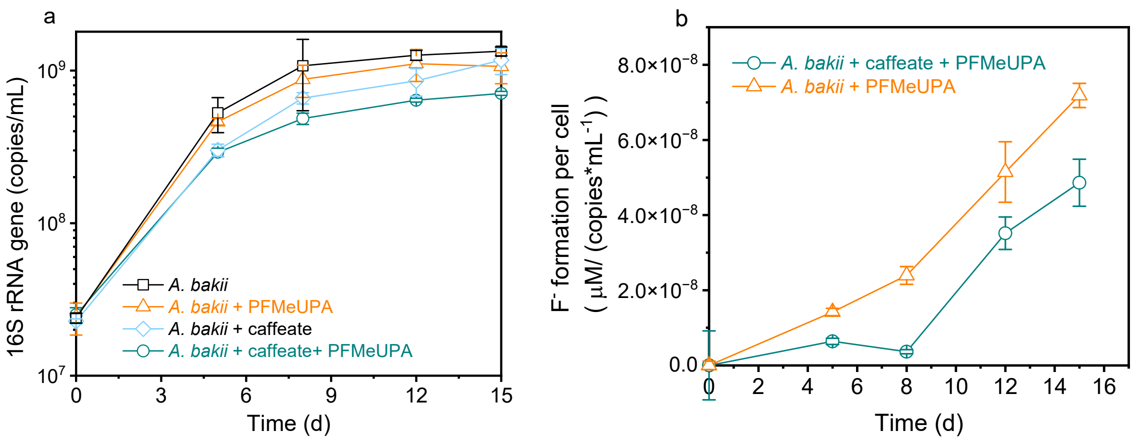


**Figure S8. Cell growth of the *A. bakii* under different substrates exposure condition (a) and normalized fluoride concentration to cell growth (b).**

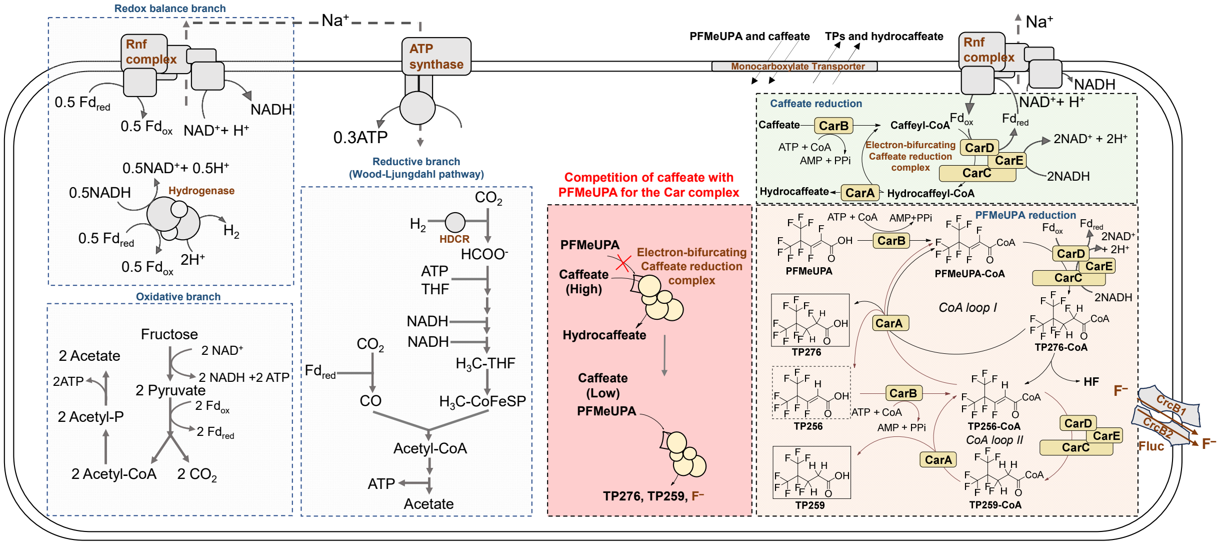


**Figure S9. Schematic illustration of the metabolic pathways in *A. bakii***. The stoichiometry of reactions in Redox balance branch, Oxidative branch, and Reductive branch (Wood-Ljungdahl pathway) are sourced from the work of Wiechmann A., Müller V., et al[8]. Modifications to caffeate reduction were adapted from Bertsch J., Müller V., et al[9]. Additionally, the PFMeUPA reduction pathways and competition models were conceptualized based on caffeate reduction pathways and *in vivo* substrates competition and modeling results (Figure 3).

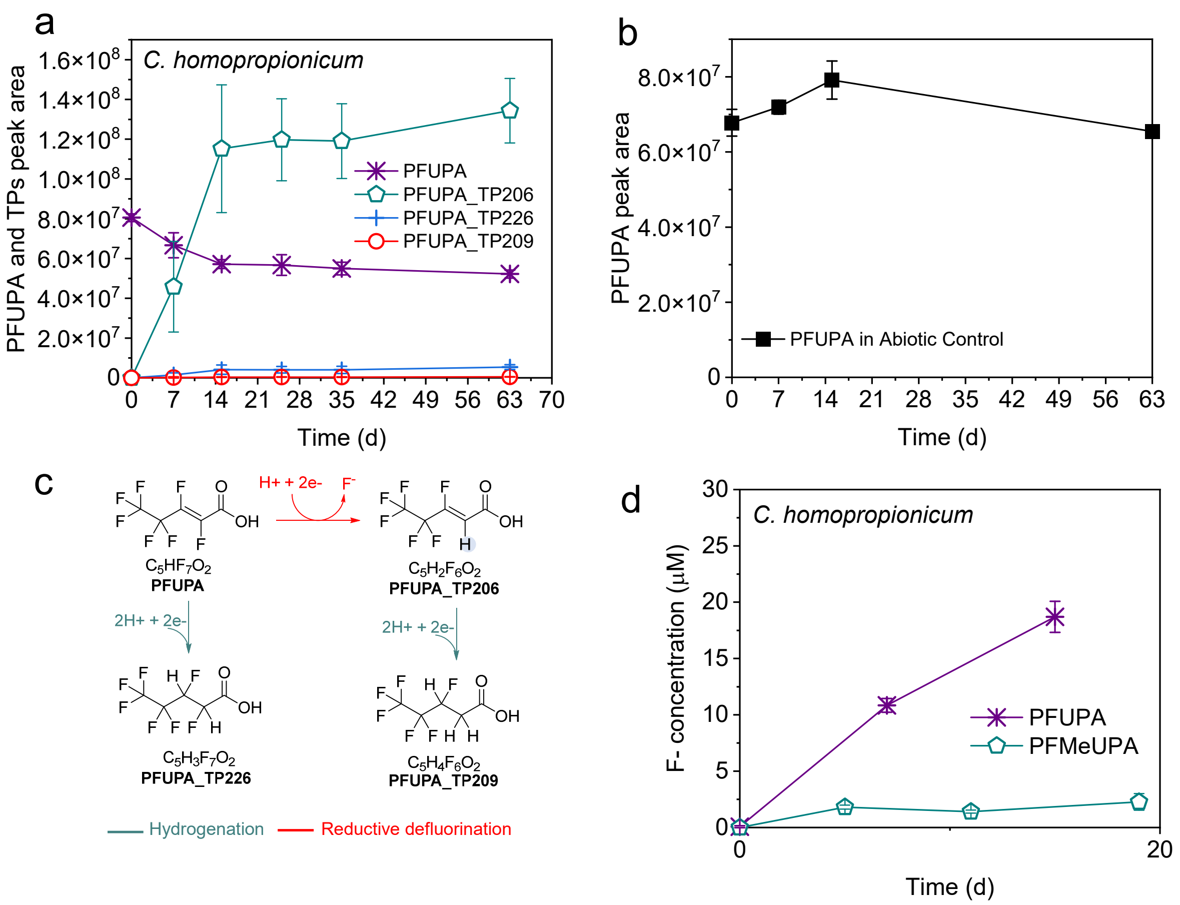


**Figure S10. Biotransformation and defluorination of a shorter chain unsaturated perfluorinated compound (PFUPA) by *Clostridium homopropionicum* DSM 5847.** a, Biotransformation of PFUPA and TPs formation. b, Abiotic degradation of PFUPA. c, Biotransformation pathways of PFUPA. d, biodefluorination of PFUPA and PFMeUPA by *C. homopropionicum.* (Note: given the impurities in PFUPA can result in abiotic defluorination[10], to determine the actual biodefluorination of PFUPA, $F^-$ concentration in panel d is calculated as:

$$\boldsymbol{PFUPA\ biodefluorination}\ (\mu\boldsymbol{M}) = \boldsymbol{F}^-_{\boldsymbol{biological\ samples}}(\mu\boldsymbol{M}) - \boldsymbol{F}^-_{\boldsymbol{abiotic\ samples}}(\mu\boldsymbol{M})$$

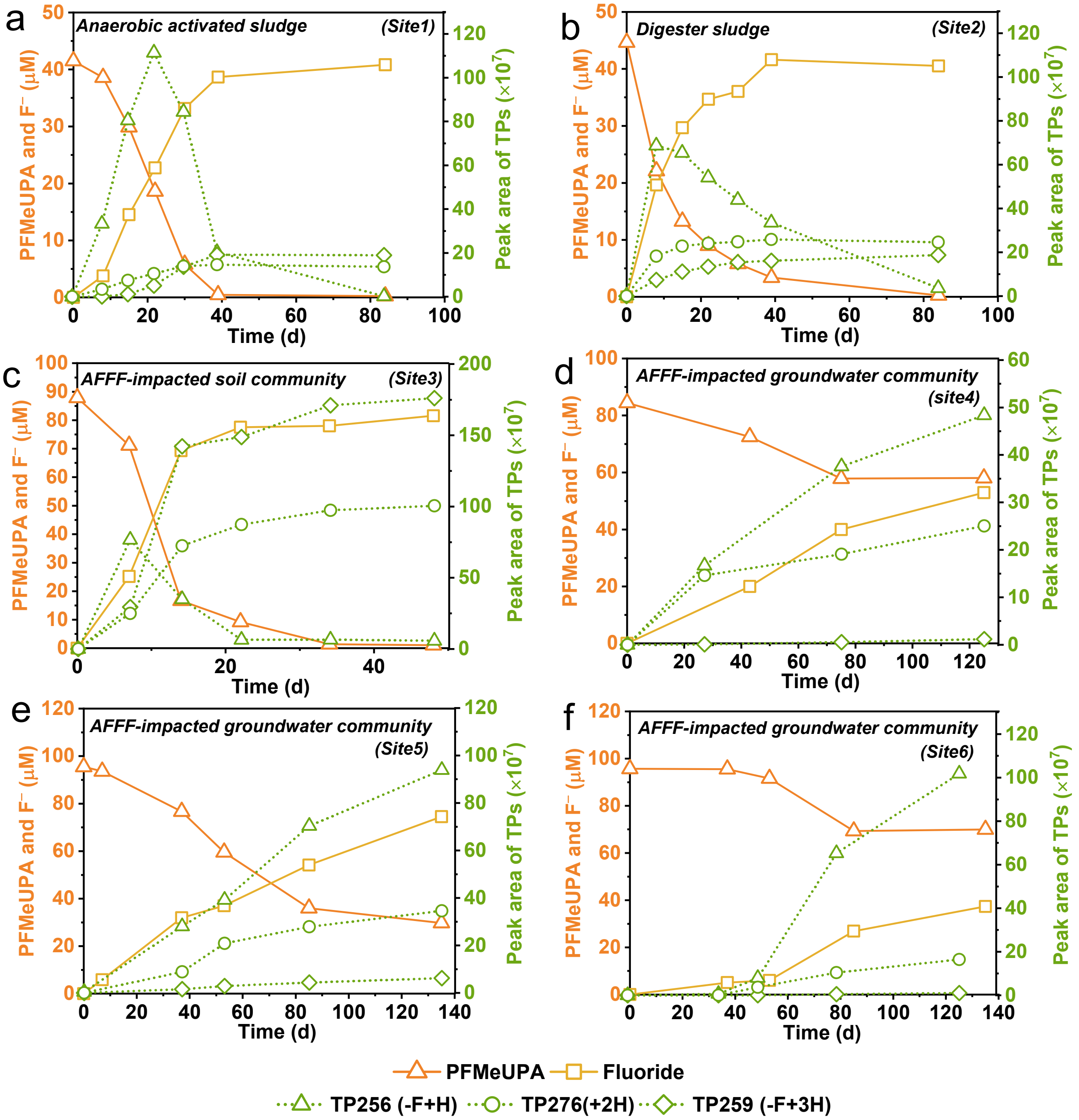


**Figure S11. Biodefluorination of PFMeUPA in environmental microbial communities taken from different sites.** a, activated sludge from a local wastewater treatment plant, which was grown in anaerobic conditions. b, digester sludge from another local wastewater treatment plant. c, soil community from AFFF-impacted site on West Coast. d-f, microbial communities in AFFF-impacted groundwater from three different sites on East and West Coasts.
